# Endotoxemia and TLR4 via tissue resident macrophages triggers anemia in mouse model of colitis

**DOI:** 10.64898/2026.03.16.712224

**Authors:** Kavita Bisht, Svetlana Shatunova, Valérie Barbier, Alnaz Husseinzoda, Ran Wang, Shrutika Mate, Ruiqing Zhong, Rabina Giri, Anna Amiss, Kylie A. Alexander, Susan M. Millard, Ingrid G. Winkler, Yoon-Kyo An, Jakob Begun, Jean-Pierre Lévesque

## Abstract

Anemia is one of the most debilitating and frequent complication of inflammatory bowel diseases (IBD) and is often treated with iron supplementation, which has limited efficacy and causes additional co-morbidities. Damaged intestinal barrier function is a hallmark of IBD and causes the translocation of endotoxins from gut bacteria into the bloodstream. A previous study in mice reported that endotoxins suppress erythropoiesis by reprogramming erythroblastic island macrophages (EBI Mφ). Here, we show that IBD patients and mice with acute colitis developed endotoxemia associated with anemia. Endotoxemia in IBD patients was negatively correlated with blood erythrocyte counts. In line with this, mice with colitis caused by drinking water containing dextrin sodium sulphate (DSS) had endotoxemia together with anemia characterized by reduced red blood cell counts, hemoglobin content and hematocrit, and reduced medullary erythropoiesis which was in part compensated by increased extramedullary erythropoiesis. As the endotoxin receptor TLR4 is expressed by CD169^+^ gut-resident macrophages as well as erythroid island macrophages in the bone marrow, we tested the hypothesis that TLR4 expressed by these CD169^+^ macrophages mediate both inflammatory colitis and anemia. Indeed, mice with a conditional deletion of the *Tlr4* gene specifically in CD169^+^ tissue-resident macrophages were protected from DSS-induced anemia, inflammation and colitis. In addition, treatment with the TLR4 inhibitor C34 abated inflammation and anemia in DSS mice. These results suggest that endotoxins leaking from the inflamed gut may play a crucial role in IBD and associated anemia via CD169^+^ tissue-resident macrophages and that drugs targeting TLR4 may protect against IBD-associated anemia.

## Introduction

The incidence of inflammatory bowel diseases (IBD) is high and continually increasing in the Americas, Europe, Asia and Oceania.^1^ Anemia is the most common comorbidity of IBD^2–4^ affecting both Crohn’s disease (CD) patients (range 10.2% – 72.7 %) and ulcerative colitis (UC) patients (range 8.8% – 66.6%).^5^ Anemia in IBD can result in low tissue oxygenation causing cardiac, metabolic and immune deficits, which together may lead to chronic fatigue, lost productivity, poor quality-of-life, absenteeism at work for weeks or months with increased rates of hospitalization, and early mortality.^3,4,6,7^

Iron deficiency anemia (IDA)^8^ and anemia of inflammation (AOI)^9^ are the two commonest causes of anemia in IBD. However, IDA and AOI often co-exist in IBD, challenging diagnosis and treatment.^10,11^ Based on the assumption that alimentary iron intestinal absorption is reduced in the inflamed gut and that intestinal bleeding increases iron and erythrocytes loss leading to iron-deficiency and poor hemoglobin synthesis, iron supplementation is considered the first-line therapy for anemia.^12–15^ However, iron supplementation fails to correct anemia in half of anemic IBD patients^4,7,16^ and cause additional co-morbidities including gastrointestinal symptoms^17^ and osteomalacia with increased bone fractures^18^, highlighting the urgent need to find better therapeutic approaches to treat IBD-associated anemia and improve clinical management of these patients. In this study, we explored the mechanisms of anemia to identify potential interventions for correcting IBD-associated anemia.

We previously reported that bacterial lipopolysaccharides (LPS; endotoxins) suppress medullary erythropoiesis in part by reprogramming erythroblastic island (EBI) macrophages (Mφ) and blocking their supportive function to complete erythroblast maturation.^19^ As intestinal barrier damage and increased intestinal permeability are characteristics of IBD,^20^ we hypothesized that the disruption of intestinal barrier function may lead to the translocation of endotoxins from gut bacteria into the bloodstream (endotoxemia) which in turn may suppresses medullary erythropoiesis, increases erythrocyte clearance in the spleen causing anemia. Indeed, 31% of CD and 17% of UC patients display endotoxemia.^21^ These prevalences are similar to that of anemia in patients with CD (29%) and UC (21%) reported in a large meta-analysis.^22^ However a functional connection between endotoxemia and anemia has not been established in IBD patients or mouse models.

In the present study, we report that endotoxemia is inversely correlated with blood erythrocyte counts in IBD patients. To investigate the mechanistic link between anemia and endotoxemia in IBD, we utilized a dextran sodium sulphate (DSS)-induced model of acute colitis in mice and observed strong association with endotoxemia and anemia. We further show that conditional deletion of the endotoxin receptor gene Toll-like receptor-4 (*Tlr4*) specifically in CD169^+^ tissue-resident macrophages (Mφ) or treatment with a small synthetic TLR4 inhibitor can significantly alleviate anemia in the DSS-induced colitis model.

## Methods

### Human subjects

We retrieved serum and clinical data from 19 non-IBD healthy controls, 38 CD and 38 UC samples stored at the Mater Hospital IBD biobank. Blood variables and tests for iron and associated parameters were performed by Mater Pathology. This Study was approved by Mater Hospital Human Research Ethics Committee (HREC/MML/89808 (V6) and Mater Research Governance (BMSSA/MRGO/89808 (V3).

### Mice

C57BL/6 mice were purchased from Ozgene (Western Australia). *Tlr4^fl/fl^* (B6(Cg)- *Tlr4*^tm1.1Karp^/J; RRID:IMSR_JAX:024872) and R26^ZsG^ (B6.Cg-*Gt(ROSA)26Sor*^tm6(CAG-ZsGreen1)Hze^/J; RRID:IMSR_JAX:007906) mice^16^ were purchased from Jackson laboratory (Bar Harbor, Maine, USA). *Siglec1^Cre^ (Siglec1*^tm2(icre)Mtka^; RRID:IMSR_RBRC06239)^17^ mice were sourced from the Riken BioResource Centre (Yokohama, Kanagawa, Japan). *Tlr4*^fl/fl^, *Siglec1^Cre^:Tlr4^fl/fl^, R26*^ZsG^ mice were bred at Translational Research Institute Biological Research Facility. *Siglec1^Cre^* mice were intercrossed with either *Tlr4*^fl/fl^ *or R26*^ZsG^ *mice* to produce either *Siglec1^Cre^:Tlr4*^fl/fl^ or *Siglec1^Cre^:R26*^ZsG^ mice. All mice used for experiments were 8-10 week-old and age- and gender-matched. All experiments were approved by The University of Queensland ethics committee (2019/AE000313, 2021/AE001177 and 2021/AE000490).

### Mouse model of colitis

Acute colitis was induced in mice by administration of 3% 40-kDa DSS (Sigma-Aldrich, Cat#42867) in drinking water for 7 days. DSS water was changed every third day. In some experiments, mice were injected intraperitoneally with 10mg/mL bromodeoxyuridine (BrdU) single dose on day 6 of DSS treatment.

### Statistical analysis

All statistical analyses were performed using GraphPad Prism (v10.4). Data are presented as mean ± SD. Statistical tests and p values are indicated in figure legend. P value <0.05 was considered significant.

Additional methods are in the Supplementary methods.

## Results

### IBD patients are anemic with increased endotoxemia and inflammation

Gut microbiota dysbiosis in IBD compromises the intestinal barrier integrity facilitating the release of endotoxins produced by gram-negative bacteria into the circulation.^20^ To assess for a possible correlation between endotoxemia and anemia in IBD subjects, we performed a retrospective study on archived non-IBD controls (Cont), CD and UC patients’ serum samples stored at Mater Hospital biobank. Patients’ characteristics including age, gender, disease severity and hematological parameters are summarized in Figure 1A. CD and UC patients had significantly lower red blood cell (RBC) counts compared to Cont, (Figure 1A-B). While other hematological parameters were not statistically altered (Figure 1A). Similarly, no statistically significant changes in inflammation marker C-reactive protein, transferrin, transferrin saturation and ferritin were observed (Figure 1A), together suggesting factors other than IDA may be contributing to anemia in these patients. We found significant increase in blood endotoxin levels in IBD patients with ∼2.7-fold increase in CD and ∼3.3-fold increase in UC patients compared to non-IBD patients (Figure 1C) consistent with previous study.^21^ Spearman correlation analysis showed a negative correlation between blood RBC counts and endotoxin levels (r = −0.3077 and p = 0.0297, Figure 1D). Most patients in this cohort had normal iron concentrations and only a few Cont (2/10), CD (6/30) and UC (4/29) patients had iron concentration below the reference range (<9.5 µmol/L, Figure 1D). Together these results suggest anemia might be functionally linked to endotoxemia and systemic inflammation. We next measured markers of inflammation in these patients. Hepcidin is a key iron transport regulator^27^ whose released is induced by inflammatory cytokine interleukin (IL)-6.^28^ Hepcidin concentration was significantly elevated in sera of CD and UC patients with a significant ∼1.2 and ∼1.3-fold increase in IL-6 concentration in CD and UC patients respectively (Figure 1F-G). Likewise, interferon (IFN)-γ, which enhances RBC clearance by splenic red pulp Mφ^29^ was significantly increased in both CD and UC patients (Figure 1H). CD and UC patients also showed significant increases in IL-1β, IL-8, IL-10, IL-12p70, IL-17a, IL-18, IL-33 and IFN-α2 blood concentrations (Supplementary Figure 1). These results indicate that endotoxin-mediated inflammation may cause anemia in IBD.

**Figure 1.**
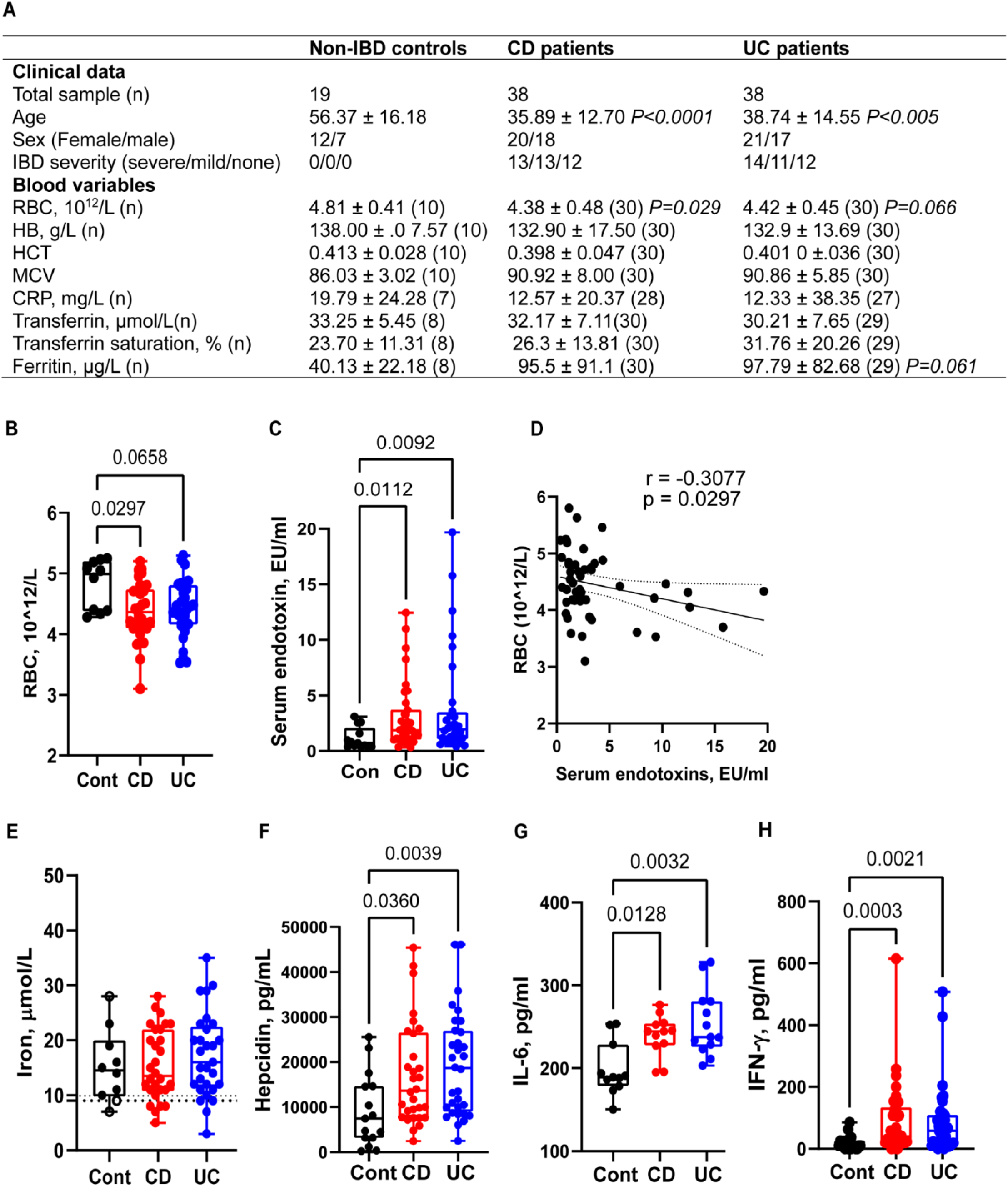
IBD patients are anemic with increased endotoxemia and inflammation. A) Characteristics of IBD patients and controls included in the study. B) Red blood cell (RBC) counts and C) serum endotoxin concentrations from controls (n=10), CD (n=30) and UC (n=29) patients. Statistical analyses were performed by Kruskal-Wallis test followed by Dunn’s post-hoc multiple comparison. C) Negative correlation between endotoxin concentration in serum and RBC in all patients was assessed by Spearman rank correlation. D) Serum iron E) hepcidin F) interleukin-6 (IL-6) and G) interferon-gamma (IFN-γ) concentrations. Statistical analyses were performed by Kruskal-Wallis test followed by Dunn’s post-hoc multiple comparison. Each dot represents a different patient. Data are presented as box and whisker plot with median, interquartile range, minimum and maximum.

### Colitis in mice causes endotoxemia and enhances extramedullary erythropoiesis

To further understand the anemia in IBD, we investigated the presence of anemia in acute colitis induced by 3% DSS in drinking water for 3-7 days in C57BL/6 mice (Figure 2A). DSS is a classical model of severe colitis in mice with features similar to UC, including intestinal crypt erosion, crypt abscess, and increased gut permeability.^30^

**Figure 2.**
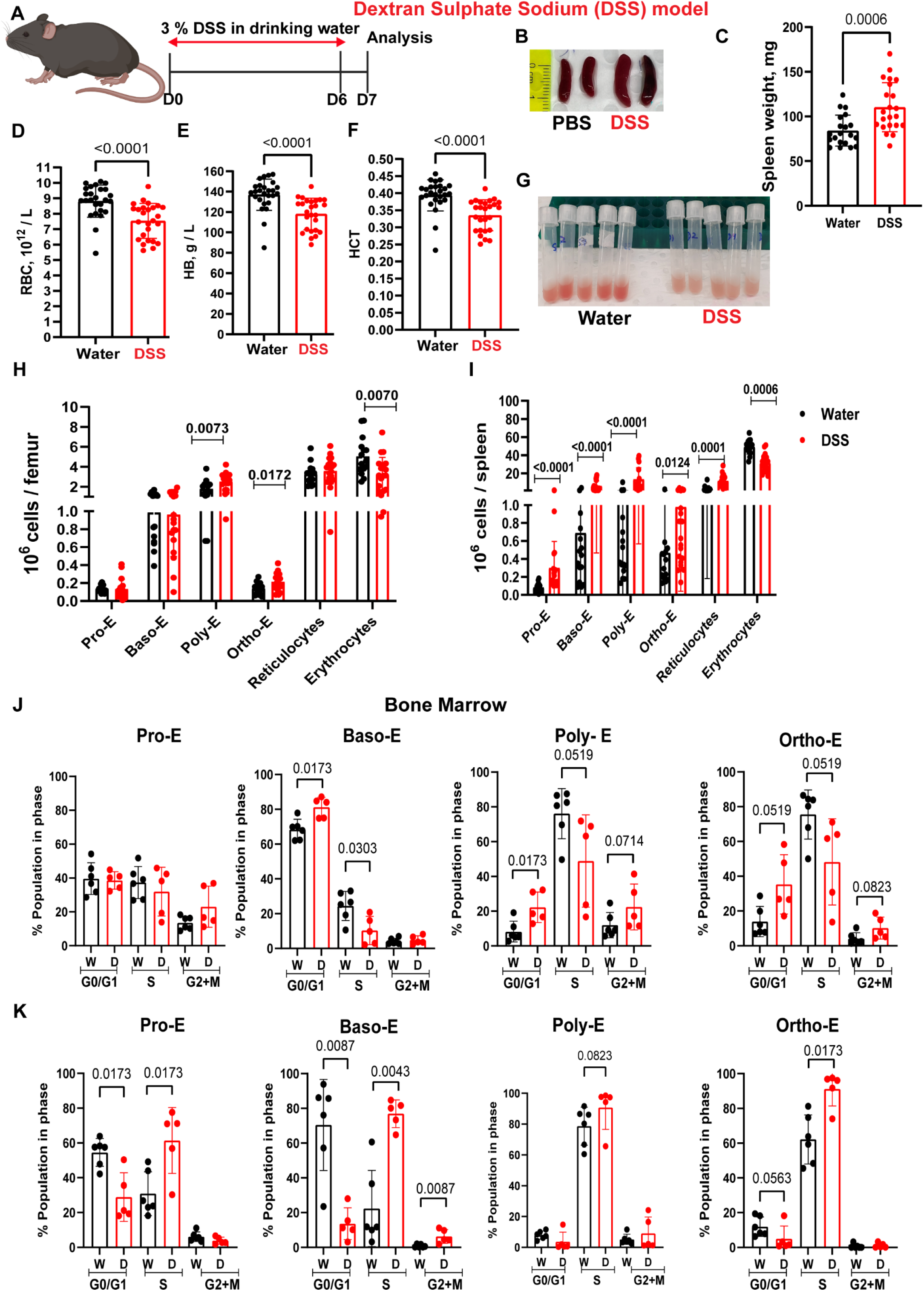
Colitis in mice causes anemia and increased extramedullary erythropoiesis. **A**) Schematic of dextran sodium sulphate (DSS)-induced colitis model. B) Photographs of mouse spleen and C) spleen weight. D) RBC, E) hemoglobin concentration (HB) and F) hematocrit (HCT) were measured with Minday hematology analyzer. Data are pooled from 6 different experiments (n= 26 mice/group total). G) Photographs of mouse femoral BM flushed into 1 mL PBS after water or DSS treatment. Note the discoloration of BM after DSS-induced colitis. H-I) Numbers of proerythroblasts (pro-E), basophilic erythroblasts (Baso-E), polychromatic erythroblasts (Poly-E), orthochromatic erythroblasts (Ortho-E), reticulocytes and erythrocyte populations in the femoral BM (H) and (I) spleen as quantified by flow cytometry. Data are pooled from 3 different experiments (n=15/water and 19/DSS groups). J-K) BrdU incorporation in proerythroblasts (pro-E), basophilic (baso-E), polychromatic (poly-E) and orthochromatic (ortho-E) was measured by flow cytometry in the BM (J) and spleen (K). n=5-6 mice/group. Dots in all charts show values for individual mice, bars represent mean ± SD and statistical analyses were performed by Mann–Whitney test.

DSS-induced colitis caused anemia in a time-dependent manner associated with spleen enlargement (Figure 2B-C) and increased extramedullary erythropoiesis in the spleen observed by day 7 of DSS treatment (Supplementary Figure 2B). Anemia was documented by significant and time-dependent decrease in RBC numbers, hemoglobin (HB) concentration and hematocrit (HCT) with a magnitude of ∼-15% for each (Figure 2D-F, Supplementary figure 2C-E). Significant increase in neutrophil number and the distribution of RBC width (RDW) along with decreased lymphocyte numbers were also observed in the blood (Supplementary Figure 3). DSS administration also induced discoloration of the BM (from red to pink; Figure 2G) and impaired erythroblast (EB) maturation at this site, with ∼1.41-fold increase in polychromatic EB (Figure 2H and population III in Supplementary Figure 4) and 37% reduction in number of erythrocytes in the BM (Figure 2H and population V in Supplementary Figure 4). The loss in BM erythropoiesis in DSS-treated mice was partially compensated by increase in extramedullary erythropoiesis in the spleen with splenomegaly (Figure 2B-C) and increased number of proerythroblasts and erythroblasts compared to control mice (Figure 2I and Supplementary Figure 4). These results establish that DSS-induced colitis is a complementary mouse model to study hematological impact of IBD.

To determine whether DSS alters erythroblast quiescence, mice were injected BrdU 24 hours before the endpoint. BrdU is a thymidine analog that is incorporated into genomic DNA during the S phase of cell division.^31^ We harvested BM and spleen and stained cells for erythroid markers, BrdU and 7AAD as described in supplementary methods. In the BM, DSS treatment increased the proportion of cells in G0/G1 phase to∼82% of basophilic, ∼22% of polychromatic and ∼ 35% of orthochromatic EBs with significantly reduced proportion or erythroblasts transiting through S-phase compared to control mice (Figure 2J). Whereas in the spleen, ∼61% of proerythroblasts, ∼77% of basophilic, ∼91% of orthochromatic EBs incorporated BrdU in S phase (Figure 2K). These results suggest that erythroblasts are proliferating less in the BM but dividing more actively in the spleen upon DSS-induced colitis.

### DSS-induced colitis causes endotoxemia, inflammation, reticulocytosis and reduces RBC lifespan in mice

We next measured blood endotoxin concentrations in DSS. DSS-treated mice had significantly elevated levels of both endotoxin (∼4.6-fold increase, Figure 3A) and inflammatory cytokines in the blood. This included a ∼9.2-fold increase in IL-6, ∼1.6-fold increase in IFN-γ (Figure 3B-C) and increases in additional cytokines in the blood (Supplementary Figure 5). This is consistent with observations in IBD patients and prior demonstrations of compromised gut barrier function in this model.^32^ When we performed correlation analysis using Spearman correlation, we found a negative correlation between blood RBC counts, HB and HCT and endotoxin levels (Figure 1D-F).

**Figure 3.**
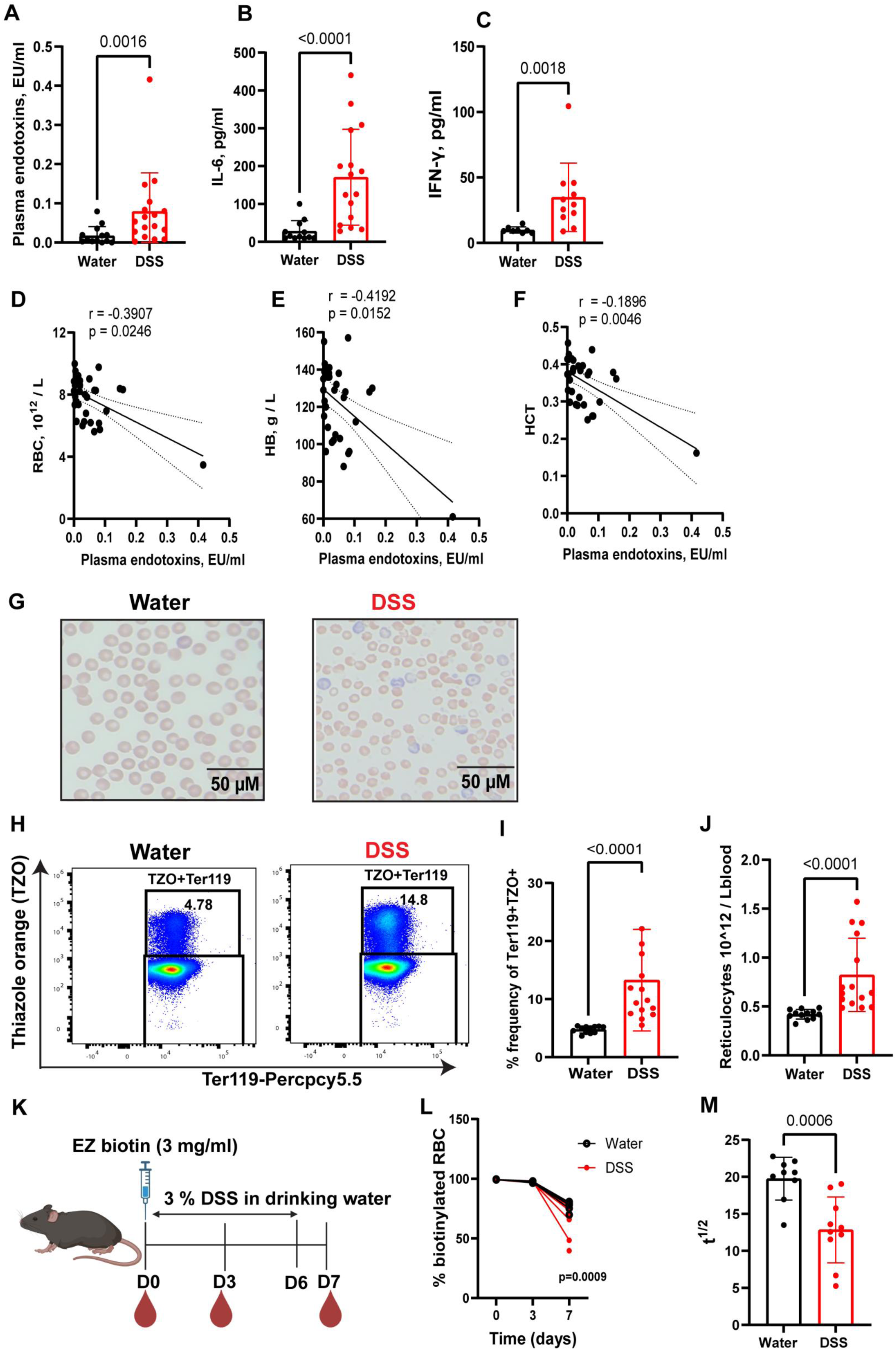
DSS-induced colitis in mice causes endotoxemia, inflammation, reticulocytosis with reduced RBC lifespan. A) Endotoxin concentration in mouse blood plasma on day 7 of DSS or plain water treatment. Data are pooled from 5 independent experiments (n=14 mice/water and 16/DSS group). B) IL-6 and C) IFN-γ concentrations in blood plasma on day 7 of DSS or plain water treatment. Data are pooled from 2 independent experiments (n=7/water and 10/DSS group). D-F) Plasma endotoxin concentration relative to RBC counts (D), hnemoblogin concentration € and hematocrit (F)., G) May-Grunwald Giemsa staining of blood smears on day 7 of treatment with DSS or plain water. Note the increase of large blue stained reticulocytes in DSS-induced colitis mice. H) Flow cytometry histogram plots showing an increase in RNA containing blood reticulocytes (TZO^+^Ter119^+^) at day 7 of DSS treatment. I) Frequency and J) numbers of reticulocytes in the blood at day 7 of DSS treatment. Data are pooled from 3 independent experiments. (n=12 mice/water and 15/DSS group). K) Schematic of EZ-biotin intravenous injection to measure RBC half-life in vivo in DSS treated mice. L) Percentage of biotinylated RBC measured by flow cytometry at indicated time points post DSS treatment. M) Calculated half-life of RBC in days. Data are pooled from 2 independent experiments (n=5-9/water and 5-10/DSS group). Statistical analyses were performed by Mann-Whitney test. Each dot is a separate mouse. Bars represent mean ± SD.

Microscopic examination of May-Grunwald Giemsa stained blood smears showed erythrocyte abnormalities, greater RBC size variation and increased number of reticulocytes in DSS-treated mice (Figure 3G), confirmed by flow cytometry analyses with significantly higher reticulocyte frequencies and counts in blood of DSS-treated mice (Figure 3H-J) in line with increased RBC width (Supplementary Figure 3). Indeed, RBC half-life measurements using an in vivo biotinylation assay or blood erythrocytes^33^ (Figure 3K) showed that DSS-treated mice exhibited accelerated loss of biotinylated RBCs with reduced RBC half-life from t_1/2_ = 19.7 ± 2.9 days in control mice to t_1/2_ = 12.8 ± 4.5 days in DSS-treated mice(Figure 3L-M).

### Mice with DSS-induced colitis have impaired iron homeostasis

Perls’ Prussian blue iron staining was performed on tissue sections as an indicator of iron stores.^34^ Perls’ Prussian blue was evident in spleen red pulp with occasional stain in the white pulp (Figure 4A-B) but undetectable in liver (data not shown) of control mice. However, splenic iron staining significantly decreased by 63% in DSS-treated mice (Figure 4A and B). We next used a colorimetric assay to quantify non-heme iron levels in spleen and liver extracts.^35^ Consistent with histological analysis, DSS-treated mice had significantly lower total tissue iron in the spleen (32% decrease, Figure 4C) but no change in liver (Figure 4D). Likewise, DSS-treated mice had significantly lower blood iron concentration and transferrin saturation (Figure 4E-F). We next assessed key components of iron homeostasis including the concentration of hepcidin in the blood and expression of the gene encoding hepcidin (*Hamp*) in the liver. Hepcidin regulates the activity of the iron exporter ferroportin (encoded by the *Slc40a1* gene), which is essential for iron cellular export. While DSS treatment had no impact on *Hamp* expression in the liver or plasma hepcidin concentration (Figure 4G-H), it significantly downregulated *Slc40a1* expression in the duodenum (86% reduction, Figure 4I) and BM (38% reduction, Figure 4J) suggesting impaired alimentary iron absorption and iron export from BM stores after DSS treatment. These results indicate that DSS-colitis-associated anemia is combination of both iron deficiency anemia^30^ and anemia of inflammation.

**Figure 4.**
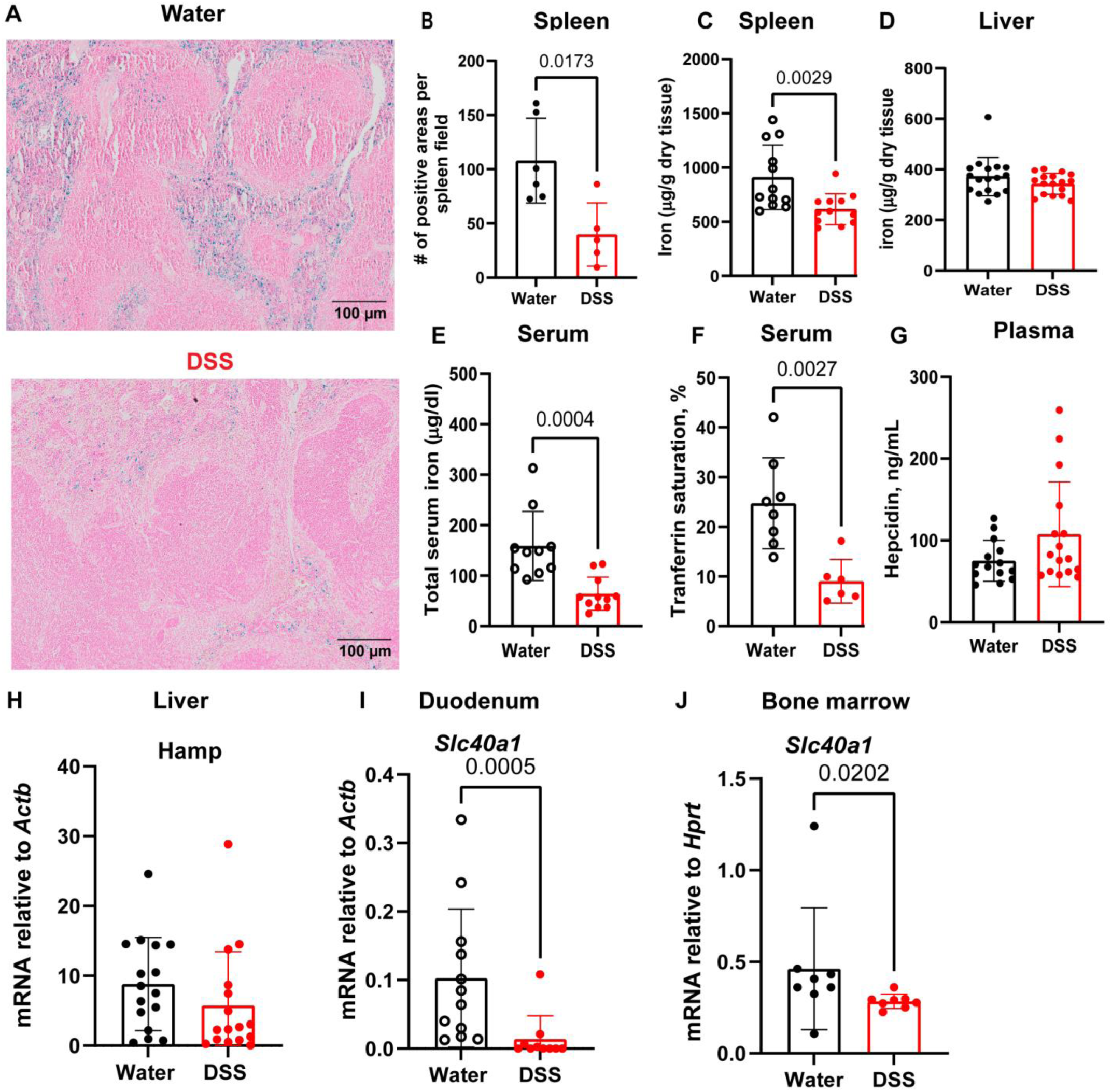
DSS-induced colitis reduces iron storage in spleen and expression of iron exporter ferroportin. A) Pearls Prussian blue staining of spleen sections from mice treated with water or DSS for 7 days (Scale bars: 100 µm). B) Positive Prussian blue staining area in the spleen was quantified with QuPath software (n=6/water and 5/DSS group). C) Colorimetric non-heme iron measurement in spleen and D) liver extracts. Data are pooled from 3 independent experiments (n=12/group). E) Serum non-heme iron, F) transferrin saturation (%) and G) plasma hepcidin concentration at day 7 of DSS or water treatment. H) Hepcidin gene (*Hamp*) expression was quantified by qRT-PCR relative to *Actb* (β-actin) in liver. Data are pooled from 2-4 independent experiments (4E and F; n=8-10/water and 6-11/DSS group and 4G; n=14-16/water and 16/DSS group). I-J) Ferroportin gene (*Slc40a1*) expression was measured in duodenum (I) and BM (J) by qRT-PCR. Data are pooled from 2- 3 different experiments (n=8-12/water and 8-10/DSS group). Statistical analyses were performed by Mann-Whitney test. Each dot represents a separate mouse. Bars represent mean ± SD.

### Tissue macrophage-specific *Tlr4* gene deletion rescues DSS-induced anemia and corrects extramedullary erythropoiesis

We have previously shown that endotoxins inhibit medullary erythropoiesis indirectly via EBI Mφ, which express the CD169 antigen (encoded by *Siglec1* gene) and TLR4.^19,36,37^ To test whether TLR4 expressed by CD169^+^ tissue-resident Mφ play a role in colitis-induced anemia, we pursued a conditional knockout strategy. *Siglec1*^Cre^ drives floxed DNA excision specifically in CD169^+^ tissue resident macrophage populations, with little activity in monocytes.^24^ *Siglec1^Cre^:Tlr4^fl/fl^*mice were generated and loss of TLR4 expression on peritoneal Mφ was confirmed by flow cytometry (Figure 5A-B) We next investigated effect of Mφ-specific *Tlr4* deletion on DSS-induced anemia. Following DSS treatment, *Siglec1^Cre^:Tlr4*^f*l/fl*^ mice were not anemic with normalized RBC, HB, HCT and reticulocyte numbers compared to *Siglec1^Cre^:Tlr4^wt/wt^* mice (Figure 5C-G). Likewise, *Siglec1^Cre^:Tlr4^fl/fl^* mice had normal spleen size and showed no extramedullary erythropoiesis with normal number of proerythroblasts and erythroblasts in the spleen in response to DSS (Figure 5H-K). Mice lacking the *Tlr4* gene specifically in CD169^+^ tissue-resident Mφ had blood endotoxin levels similar to non-DSS treated mice (Figure 5L). Systemic inflammation upon DSS treatment was blunted in *Siglec1^Cre^:Tlr4^fl/fl^* mice with a 53% decrease in IL-6 blood concentration although it remained well above basal levels (Figure 5M). *Siglec1^Cre^:Tlr4^fl/fl^* mice showed an increase trend for Perls’ Prussian blue iron staining in the spleen (Figure 5N-O, P=0.0996) suggesting that TLR4 via CD169^+^ tissue-resident Mφ reduces splenic iron stores.

**Figure 5.**
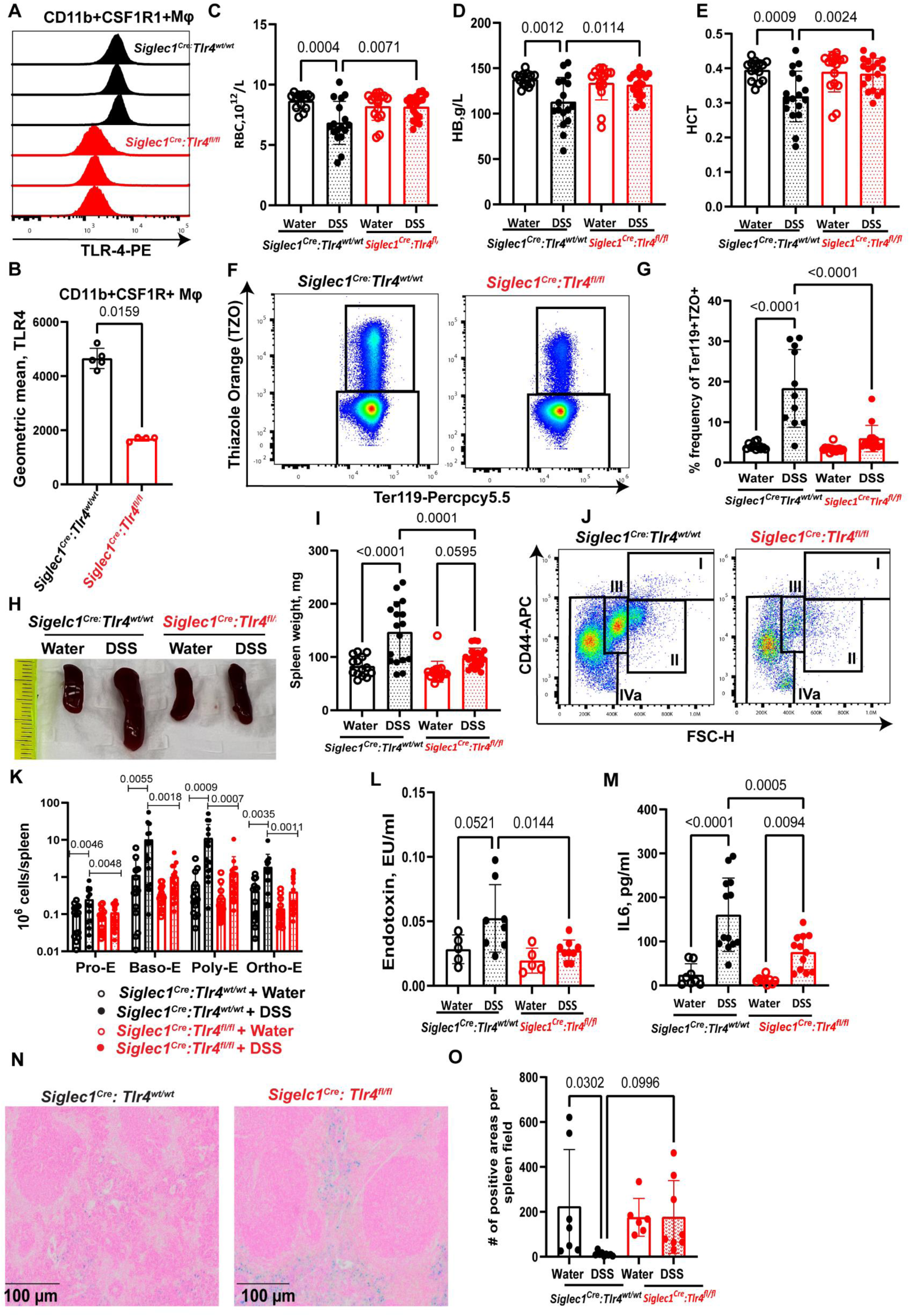
Conditional deletion of the *Tlr4* gene in CD169^+^ tissue-resident macrophages (Mφ) reduces DSS-induced anemia, endotoxemia and systemic inflammation. A) Flow cytometry histograms of TLR4 expression on CD11b^+^CSF1R^+^Ly6G^-^CD169^+^ peritoneal Mφ in *Siglec1^Cre^:Tlr4^fl/fl^*mice (red) and control *Siglec1^Cre^:Tlr4^WT/WT^* mice (black). B) Frequency of TLR4^+^ cells gated on CD11b^+^CSF1R^+^Ly6G^-^CD169^+^ peritoneal Mφ.. Statistical analyses were performed by Mann–Whitney test. Each dot is a separate mouse. Bars represent mean ± SD (n=5/ *Siglec1^Cre^:Tlr4^WT/WT^* and n=4/ *Siglec1^Cre^:Tlr4^fl/fl^*group). C) RBC counts, D) HB concentrations and E) HCT in the blood after 7 days treatment with DSS or plain water. Data are pooled from 4 independent experiments (n=14/ *Siglec1^Cre^:Tlr4^WT/WT^*+ water, n=17/ *Siglec1^Cre^:Tlr4^WT/WT^* +DSS, n=16/ *Siglec1^Cre^:Tlr4^fl/fl^* + water and n=18/ *Siglec1^Cre^:Tlr4^fl/fl^*+ DSS group). F) Flow cytometry plots showing lower TZO^+^Ter119^+^ blood reticulocyte frequency in *Siglec1^Cre^:Tlr4^fl/fl^* mice compared to control *Siglec1^Cre^:Tlr4^WT/WT^* mice at day 7 of DSS treatment. G) Quantification of reticulocytes in the blood at day 7 of DSS or water treatment in *Siglec1^Cre^:Tlr4^fl/fl^* mice (red) and control *Siglec1^Cre^:Tlr4^WT/WT^* mice (black). Data are pooled from 3 independent experiments (n=11/*Siglec1^Cre^:Tlr4^WT/WT^* + water, n=11/*Siglec1^Cre^:Tlr4^WT/WT^* + DSS, n=14/ *Siglec1^Cre^:Tlr4^fl/fl^* ± DSS group). H) Photographs of mouse spleens and I) spleen weights. J) Flow cytometry plots showing changes in numbers of proerythroblast and erythroblast subsets (I : Pro-E, II : Baso-E, III; Poly-E, IVa : Ortho-E) in DSS treated *Siglec1^Cre^:Tlr4^WT/WT^* and *Siglec1^Cre^:Tlr4^fl/fl^* mice. K) Number of proerythroblasts and erythroblasts per spleen quantified by flow cytometry (subsets described in more detail Supplementary Figure 4). Data are pooled from 4 independent experiments (n=14/ *Siglec1^Cre^:Tlr4^WT/WT^*+ water, n=17/ *Siglec1^Cre^:Tlr4^WT/WT^* +DSS, n=16/ *Siglec1^Cre^:Tlr4^fl/fl^* + water and n=20/ *Siglec1^Cre^:Tlr4^fl/fl^* + DSS group). L) Concentration of endotoxin and M) IL-6 in blood. N) Pearls Prussian blue staining of spleen sections from *Siglec1^Cre^:Tlr4*^WT/WT^ and *Siglec1^Cre^:Tlr4*^fl/fl^ mice treated for 7 days with DSS. O) Positive Prussian blue staining area in the spleen was quantified with QuPath software. Data are pooled from 2-3 independent experiments (n=5-8/ *Siglec1^Cre^:Tlr4^WT/WT^* + water, n=8-13/ *Siglec1^Cre^:Tlr4^WT/WT^* +DSS, n=5-9/ *Siglec1^Cre^:Tlr4^fl/fl^* + water and n=8-12/ *Siglec1^Cre^:Tlr4^fl/fl^* + DSS group). Each dot is a separate mouse. Bars represent mean ± SD. Statistical analyses were performed by One-Way ANOVA with Sidak’s multiple comparison test.

We next tested whether *Tlr4* deletion in CD169^+^ tissue-resident Mφ resolved LPS-induced changes on peripheral blood counts and medullary and extramedullary erythropoiesis. Mice were injected with 2.5 mg/kg LPS once daily for two days as described previously.^19,38^ As expected LPS caused monocytosis (3.2-fold)), neutrophilia (4.9-fold), eosinophilia (3.1-fold), lymphopenia (41% reduction) and thrombocytopenia (69% reduction) in the blood (Supplementary Figure 6). However, there was no change in RBC, HB concentration, HCT, MCV, MCH, RDW-CV or RDW-SD (Supplementary Figure 6) in agreement with previous reports.^19,38^ LPS-induced monocytosis, neutrophilia, eosinophilia, lymphopenia and thrombocytopenia were resolved in *Siglec1^Cre^:Tlr4^fl/fl^* with 68% decrease in monocytes, 12% increase in neutrophil, 49% increase in eosinophil counts and 57% increase in lymphocytes and 152% increase in platelet numbers. Noticeably, the BM from *Siglec1^Cre^:Tlr4^fl/fl^* mice were not whitened unlike their WT counterparts and have normal BM cellularity and spleen weight (Supplementary Figure 6N-Q). Flow cytometry confirmed that LPS treatment did not suppress proerythroblasts, basophilic, polychromatic and orthochromatic erythroblasts in *Siglec1^Cre^:Tlr4^fl/fl^* mice (Supplementary Figure 6R), establishing that LPS suppresses medullary erythropoiesis via CD169^+^ tissue resident Mφ and TLR4. However, we did not see any significant increase in extramedullary erythropoiesis in both *Siglec1^Cre^:Tlr4^WTWTl^* and *Siglec1^Cre^:Tlr4^fl/fl^*mice and spleen weights were not significantly increased in response to LPS (Supplementary Figure 6O,Q).

We then investigated the effect of *Tlr4* deletion on colitis symptoms and colon inflammation. *Siglec1^Cre^* efficacy at driving flox recombination in Mφ of the colon was assessed using *Siglec1^Cre^:R26^ZsG^*reporter mice in which ZsGreen fluorescent reporter is induced upon Cre-mediated recombination of an upstream STOP codon cassette.^39^ ZsGreen fluorescence was primarily induced in CD169^+^ and IBA1^+^ Mφ in the colon (Figure 6A) confirming both CD169 expression by tissue-resident Mφ in the gut and suitability of the *Siglec1*^Cre^ driver. *Siglec1^Cre^:Tlr4^fl/fl^* mice did not lose body weight following DSS treatment with improved colon length and colon weight ratio (Figure 6B-D). Histology confirmed that CD169^+^ Mφ-specific *Tlr4* deletion also partially restored colon villi morphology and colon histology scores, and significantly reduced IL-6 produced in cultures of colon explants from these mice (Figure 6E-G).

**Figure 6.**
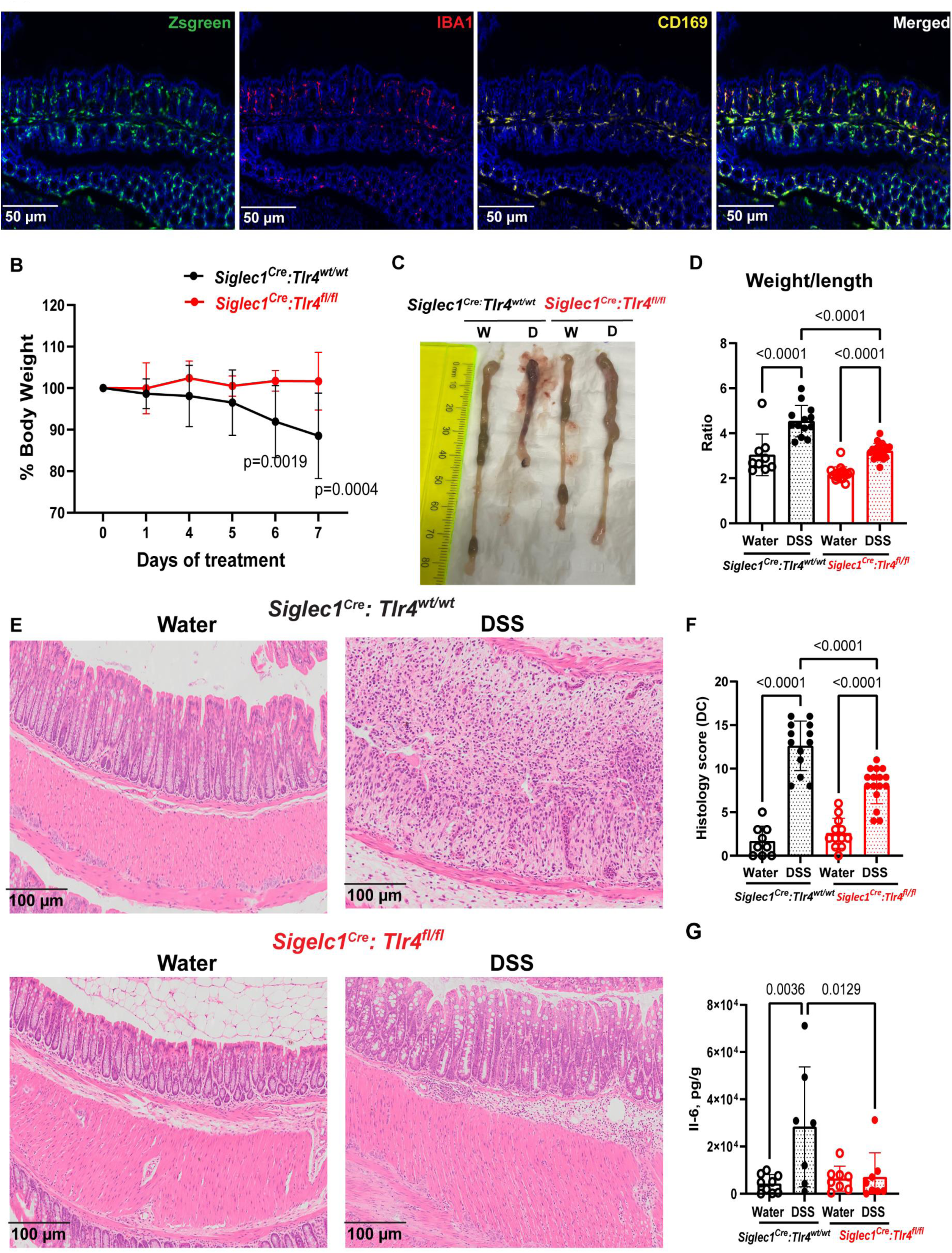
Conditional deletion of *Tlr4* gene in tissue-resident Mφ reduces DSS-induced colitis. A) Representative immunohistofluorescence images of colon sections from *Siglec1^Cre^:R26^ZsG^*reporter mice. From left to right: fluorescence of ZsGreen reporter, IBA1 (red), CD169 (yellow) tissue-resident Mφ antigens and overlayed signals (n=3). B) Kinetics of relative mouse body weight changes through DSS treatment in *Siglec1^Cre^:Tlr4*^fl/fl^ (red) and *Siglec1^Cre^:Tlr4*^WT/WT^ mice (black). Data are pooled from 4 independent experiments (n=13/*Siglec1^Cre^:Tlr4^WT/WT^*+DSS, n=20/*Siglec1^Cre^:Tlr4^fl/fl^* + DSS group). Statistical analyses were performed by Two-Way ANOVA with Sidak’s multiple comparison test. Each dot represents mean ± SD. C) Photograph of colon length of *Siglec1^Cre^:Tlr4*^WT/WT^ and *Siglec1^Cre^:Tlr4*^fl/fl^ mice treated for 7 days with DSS (D) or water (W). D) Colon weights and lengths were recorded at the end point and weight over length ratio calculated for each mouse. Data are pooled from 4 independent experiments (n=14/*Siglec1^Cre^:Tlr4^WT/WT^* + water, n=17/*Siglec1^Cre^:Tlr4^WT/WT^*+DSS, n=16/*Siglec1^Cre^:Tlr4^fl/fl^* + water and n=20/*Siglec1^Cre^:Tlr4^fl/fl^* + DSS group). Statistical analyses were performed by One-Way ANOVA with Sidak’s multiple comparison test. Each dot is a separate mouse. Bars represent mean ± SD. E) Representative hematoxylin and eosin staining of mouse colon sections and F) quantification of histology scores of distal colon (DC) in *Siglec1^Cre^:Tlr4*^WT/WT^ and *Siglec1^Cre^:Tlr4*^fl/fl^ mice treated for 7 days with water or DSS. Data are pooled from 3 independent experiments (n=9/*Siglec1^Cre^:Tlr4^WT/WT^* + water, n=13/*Siglec1^Cre^:Tlr4^WT/WT^*+DSS, n=12/*Siglec1^Cre^:Tlr4^fl/fl^* + water and n=16/*Siglec1^Cre^:Tlr4^fl/fl^* + DSS group). Statistical analyses were performed by One-Way ANOVA with Sidak’s multiple comparison test. G) IL-6 concentration in culture supernatants of colon explants isolated from *Siglec1^Cre^:Tlr4*^WT/WT^ (black) and *Siglec1^Cre^:Tlr4*^fl/fl^ (red) mice treated for 7 days with water or 3% DSS. Data are from two independent experiments (n=7-9/group). Statistical significances were calculated by One-Way ANOVA with Sidak’s multiple comparison test. Each dot represents result from colon explant from a separate mouse. Bars represent mean ± SD.

### TLR-4 antagonism rescues DSS-induced anemia and corrects extramedullary erythropoiesis but does not ameliorate DSS-induced colitis

The interaction between endotoxin (LPS) and its receptor TLR4 can be blocked by drugs that bind either endotoxin (such as colistin) or TLR4 (C34). Colistin (polymyxin E) is an antibiotic that binds to and neutralizes LPS from gram-negative bacterial cell wall.^40,41^ I*n vitro* cultures of mouse BM-derived monocytes (BMDMs) were used to validate drug efficacy. BMDMs were either treated with LPS ± colistin or LPS ± C34. Both colistin and C34 attenuated the release of IL-6, IFN-γ and other inflammatory cytokines in response to BMDM stimulation with LPS *in vitro* (Supplementary Figure 7) demonstrating these drugs are effective in blocking LPS/TLR4 interaction and signaling in vitro. The impact of these drugs on the DSS-colitis model was subsequently evaluated. To counter the potential nephrotoxicity of systemic use of colistin treatment,^42^ mice receiving colistin also received L-methionine which has demonstrated nephroprotective effects.^25^ Parenteral colistin administration did not significantly rescue DSS-induced anemia, colitis or body weight loss, which was even worse possibly due to colistin toxicity, although IL-6 blood concentration was significantly reduced (Supplementary Figure 8).

The efficacy of the small TLR4 antagonist C34, at resolving DSS-induced anemia was assessed using either oral or parenteral administration. We first administered C34 daily with DSS to mice via oral route as previously reported^43^ but this regimen had no beneficial effect on DSS-induced anemia or colitis (Supplementary Figure 9). However, when C34 was administered parenterally, via intraperitoneal injections, anemia was remarkably improved with significantly higher RBC, HB, HCT, reduced reticulocyte blood counts compared to DSS with vehicle (Figure 7B-F). Additionally, intraperitoneal C34 treatment improved medullary and extramedullary erythropoiesis. Erythrocyte numbers in the BM, spleen size, proerythroblast and erythroblast numbers in the spleen were all normalized compared to DSS-alone (Figure 7G-L). Intraperitoneal C34 treated mice demonstrated reduced endotoxemia, with blood concentrations of both endotoxin and IL-6 blood significantly lower than mice in the DSS-alone group (Figure 7M-N). Intraperitoneal C34 significantly improved Perls’ Prussian blue iron staining in the spleen with ∼3.4-fold increase compared to DSS alone (Figure 7O-P). Unexpectedly, intraperitoneal C34 treatment did not improve DSS-induced colitis severity and body weight loss (Supplementary Figure 10). Together, these results suggest that TLR4 plays an important role in DSS-induced anemia and systemic administration of drugs blocking TLR4 may be of therapeutic benefit to treat IBD-induced anemia.

**Figure 7.**
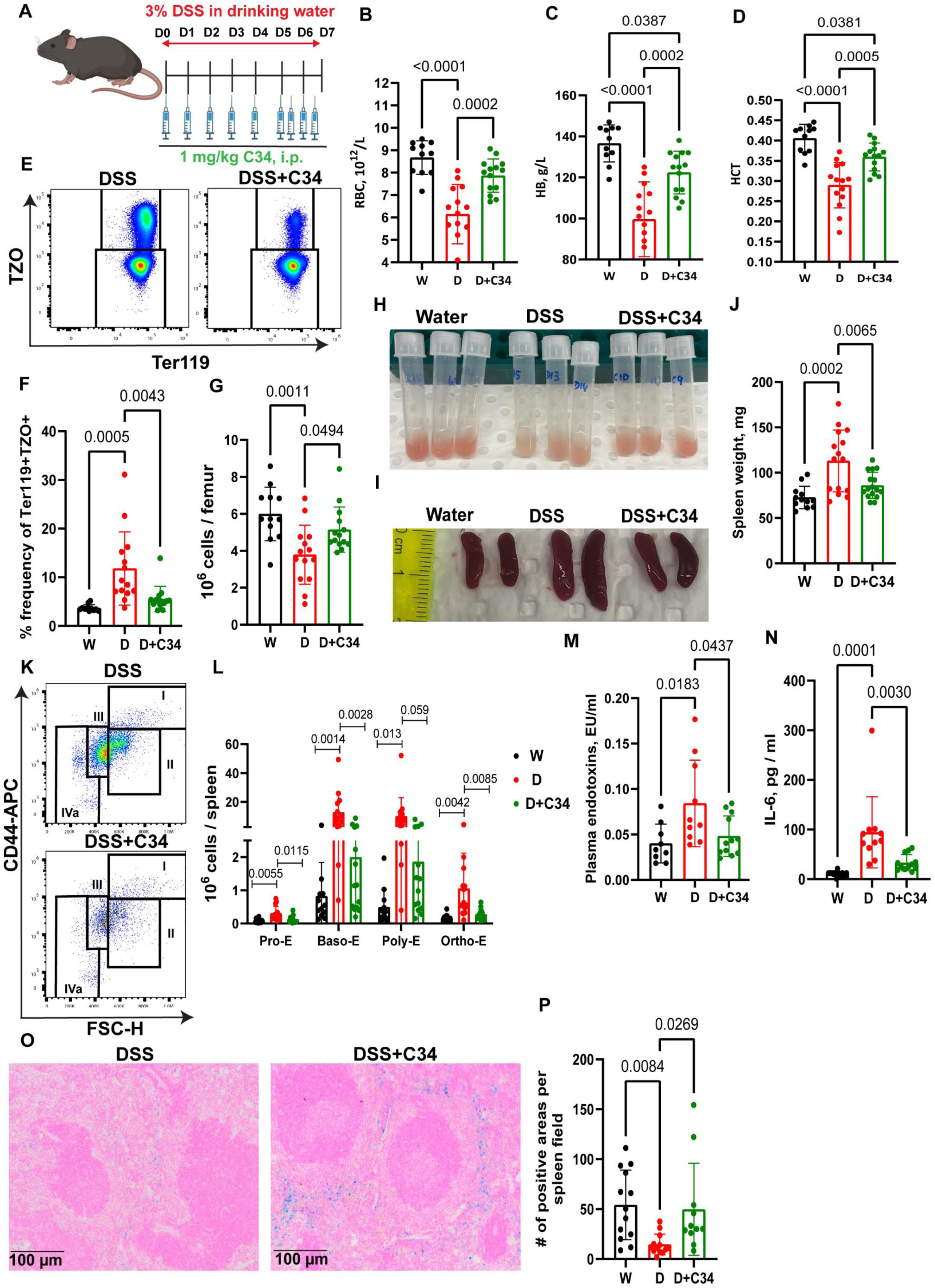
Intraperitoneal administration of TLR4 inhibitor C34 corrects DSS-induced anemia and endotoxemia. A) Schematic of C34 and DSS treatments. B) RBC counts, C) Hb concentration and D) hematocrit were measured in whole blood at day 7 of treatment with water (W), DSS (D), or DSS and C34 (D+C34). E) Flow cytometry plots and F) frequency of blood reticulocytes (TZO^+^Ter119^+^) at day 7 of treatment. G) Number of erythrocytes in the BM at day 7 of treatment. H) Photographs of mouse femoral BM flushed into 1 mL PBS and I) spleens. J) Spleen weights at day 7 of treatment. K) Flow cytometry plots showing changes in number of splenic proerythroblasts and erythroblasts (I : Pro-E, II : Baso-E, III : Poly-E, Iva : Ortho-E) in DSS±C34 groups. L) Number of proerythroblasts and erythroblasts in the spleen were quantified by flow cytometry. Data are pooled from 3 independent experiments (n=11-12/W, n=14/D and D+C34 group). Statistical analyses were performed by One-Way ANOVA with Sidak’s multiple comparison test. Each dot is a separate mouse. Bars represent mean ± SD. M) Plasma endotoxin and N) IL-6 concentrations at day 7 of treatment. O) Pearls Prussian blue staining of spleen sections from DSS±C34 group. P) Positive Prussian blue staining area in the spleen was quantified with QuPath software. Data are pooled from 2 independent experiments (n=8-13/W, 10-13/D and 11-12/D+C34 group). Statistical analyses were performed by One-Way ANOVA with Sidak’s multiple comparison test. Each dot is a separate mouse. Bars represent mean ± SD.

### TLR4 deletion in CD169^+^ tissue resident Mφ or antagonism by TLR-4-IN-C34 reduces infiltration of inflammatory monocytes

Our data suggest that TLR-4 via CD169^+^ tissue-resident Mφ promotes anemia in DSS-induced colitis and treatment with small molecule inhibitor C34 partially resolves DSS-induced anemia, suggesting that CD169^+^ tissue resident Mφ are critical in DSS-induced anemia. However, analysis of tissue resident Mφ in hematopoietic tissue is hampered by current single cell preparation as Mφ fragment during single cell suspension preparation and die.^44^ Therefore, we investigated the effect of DSS on the inflammatory activation state of different monocyte populations in blood, BM and spleen in mice lacking *Tlr4* gene in tissue resident Mφ or in C57BL/6 (WT) mice following TLR4 antagonism with C34 (gating strategy in Supplementary Figure 11). Treatment of *Siglec1^Cre^:Tlr4^wt/wt^* mice or WT mice with DSS significantly increased the number of CSF1R^+^CD64^+^ activated monocytes and LY6C^high^CCR2^+^CX3CR1^low^ inflammatory monocytes^45^ in the blood, BM and spleen (Figure 8A-F) whereas LY6C^low^CCR2^low^CX3CR1^high^ patrolling monocytes^46^ remained unchanged after DSS treatment in blood and BM of *Siglec1^Cre^:Tlr4^fl/fl^* or WT mice (Figure 8A-B, D-E). However, DSS remarkably increased the number of LY6C^low^CCR2^low^CX3CR1^high^ patrolling monocytes in the spleen of *Siglec1^Cre^:Tlr4^wt/wt^* or WT mice with ∼2-3-fold increase for each.

**Figure 8.**
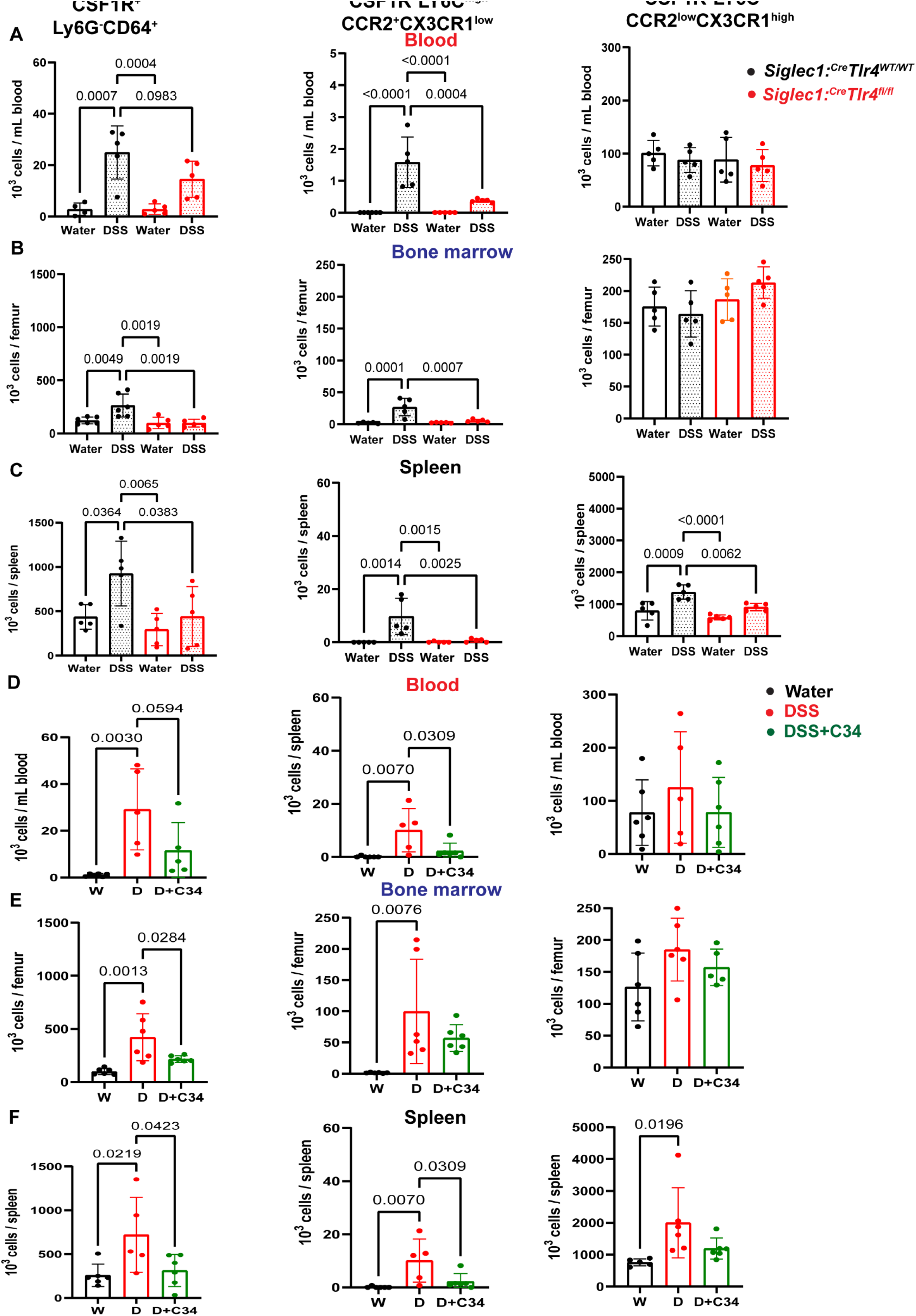
TLR4 deletion in CD169^+^ tissue resident Mφ or antagonism with TLR-4-IN-C34 reduces infiltration of inflammatory monocytes. A-F) Number of CD11b^+^CSF1R^+^Ly6G^-^CD64^+^ monocytes, CD11b^+^CSF1R^+^Ly6G^-^LY6C^high^CCR2^+^CX3CR1^low^ inflammatory monocytes and CD11b^+^CSF1R^+^Ly6G^-^LY6C^low^CCR2^low^CX3CR1^high^ patrolling monocytes were measured by flow cytometry (gating strategy in Supplementary Figure 11) in whole blood (A, D), BM (B, E) and spleen (C, F) in *Siglec1^Cre^:Tlr4^fl/fl^* mice (red) and control *Siglec1^Cre^:Tlr4^WT/WT^*mice (black) (A-C) or C5BL/6 mice at day 7 of treatment with water (W), DSS (D), or DSS with C34 (D+C34) (D-F). N=4-6/ group). Statistical analyses were performed by One-Way ANOVA with Sidak’s multiple comparison test. Each dot is a separate mouse. Bars represent mean ± SD.

DSS-induced increase in inflammatory monocytes was significantly attenuated in *Siglec1^Cre^:Tlr4^fl/fl^* mice with 40-50% decrease in CSF1R^+^CD64^+^ activated monocytes, 76-93% decrease in LY6C^high^CCR2^+^CX3CR1^low^ inflammatory monocytes in the blood, BM and spleen compared to *Siglec1^Cre^:Tlr4^wt/wt^* mice (Figure 8A-C). Likewise, treatment with C34 significantly abated DSS-induced expression of CSF1R^+^CD64^+^ activated monocytes, and inflammatory monocytes in the blood, BM and spleen by 40%-80% (Figure 8D-E). Although C34 treatment did not decrease numbers of patrolling monocytes in the spleen, *Siglec1^Cre^:Tlr4^fl/fl^* mice had significantly reduced patrolling monocytes in the spleen in response to DSS. These results suggest that endotoxin-TLR4 interaction on CD169^+^ tissue resident Mφ is required for colitis-induced activation and recruitment of inflammatory monocytes in the blood, BM and spleen.

## Discussion

In IBD, the damaged inflamed gut epithelial barrier can lead to a leaky gut with translocation of intestinal microbial products such as endotoxins into the circulation.^20,21^ Here, we identified that endotoxemia and TLR4 are important drivers of anemia in colitis in mice and discovered a positive correlation between anemia and endotoxemia in IBD patients. We also demonstrated that deletion of the endotoxin receptor *Tlr4* gene on CD169^+^ tissue-resident macrophage or intraperitoneal administration of a small TLR4 antagonist abates colitis-induced anemia in mice treated with DSS. The role of endotoxemia in colitis-induced anemia has not been extensively studied, despite prior observations that DSS-induced colitis in mice decreased HB and HCT,^47^ and that exogenous LPS further exacerbated DSS-induced colitis.^48^ To expand previous findings, we examined the effect of colitis on erythropoiesis, demonstrating that colitis worsened BM erythropoiesis, and induced splenomegaly and increased extramedullary erythropoiesis (Figure 2). Similar to IBD patients and previous studies, mice challenged with DSS also displayed significant endotoxemia and anemia in blood.^47,48^

Pro-inflammatory cytokines such as IL-6 and IFN-γ have been well documented as key candidates in anemia and directly affect erythropoiesis by reducing BM erythroid progenitor cells and shortening erythrocyte lifespan.^2,49–51^ As expected, IL-6, IFN-γ and other pro-inflammatory cytokines were significantly elevated in IBD patients and in mice after DSS and TTX-induced colitis. A shortened erythrocyte lifespan has been observed in inflammatory setting and has been attributed to increased erythrophagocytosis via hepatic and splenic red pulp Mφ.^52–54^ In line with this, we find that DSS led to abnormal erythrocyte morphology and numbers with increased reticulocyte counts together with shortened erythrocyte half-life (Figure 3). Iron homeostasis was also perturbed in DSS treated mice with low spleen and serum iron and reduced ferroportin expression in duodenum and BM suggesting impaired iron uptake and export. In a previous study, Basseri et al^55^ reported that the iron-regulatory hormone hepcidin is positively correlated with AOI and IL-6 levels in CD patients, suggesting that IL-6-induced hepcidin mediates anemia in IBD. Although we found significantly increased levels of hepcidin in CD and UC patients in accordance with previous study,^55^ we did not detect alterations in hepcidin expression or concentration after DSS-induced colitis in mice. This is consistent with a previous study by Gotardo et al^56^ with no changes in serum and liver hepcidin in a trinitrobenzene sulfonic acid (TNBS)-induced colitis model, suggesting that colitis-induced anemia in rodents is caused by combination of iron deficiency and AOI due to chronic gastrointestinal blood loss together with systemic inflammation.^8^ Our results also indicate that the spleen may recycle more iron in DSS-treated mice to meet the demand of developing EBs and facilitate extramedullary erythropoiesis.

It is known that TLRs play a crucial role in mucosal innate immune regulation.^57^ TLR4, a pattern recognition receptor for bacterial LPS and endogenous damage-associated molecular patterns (DAMPs) triggers a cascade of immune and inflammatory responses by activating two signaling pathways downstream of two adaptor proteins carrying TIR (Toll-interleukin receptor) domains, namely TRIF/TICAM1 and MyD88.^58^ Previous studies suggest a nonimmune function of TLR4 in IBD in the maintenance of epithelial homeostasis and protection from epithelial injury. Mice with germinal deletion of the *Tlr4* gene displayed severe mortality and morbidity in response to oral DSS.^48,57^ Although TLR4 deficiency abated DSS-induced secretion of pro-inflammatory cytokines, these mice were more prone to anemia,^48^ suggesting TLR4 expression plays a crucial role in gut injury and repair process. It has been shown previously that bacterial endotoxins suppress erythropoiesis via TLR4 through reprogramming of EBI Mφ which express TLR4 unlike EBs which don’t.^19^ In this study we focused on endotoxin-TLR4 signaling to gain insights into the role of this pathway on colitis-associated anemia. Since mice germinally defective for *Tlr4* gene (*Tlr4^-/-^)* are more susceptible to DSS-induced colitis,^48,57^ we deleted the *Tlr4* gene specifically in CD169^+^ tissue-resident Mφ (*Siglec1^Cre^:Tlr4^fl/fl^*mice), which include both EBI Mφ^19,36,37^ and gut-resident Mφ (Figure 6A). Of note, CD169 is expressed by only 0.5% of blood monocytes^24^. This specific deletion of *Tlr4* in CD169^+^ tissue-resident Mφ (Figure 5 A-B) allowed us to examine the effect of tissue Mφ on DSS-induced anemia. *Siglec1^Cre^:Tlr4^fl/fl^* mice were not anemic and had normalized blood counts in response to oral DSS, with neither endotoxemia nor extramedullary erythropoiesis, and systemic inflammation was blunted. *Siglec1^Cre^:Tlr4^fl/fl^*mice were also less susceptible to DSS-induced colon injury and local inflammation, suggesting that TLR4 on tissue-resident Mφ exacerbate DSS-induced colitis and anemia. Having shown protective effects of CD169^+^ tissue-resident Mφ specific *Tlr4* deletion, we next examined the effects of LPS and TLR4 inhibitors on DSS-induced anemia and colitis. Despite its LPS-neutralizing activity^41^ and strong anti-inflammatory effects on LPS-stimulated BMDMs, intraperitoneal colistin administration had no protective effect against DSS-induced anemia or colitis. This suggests that sterile inflammation driven by DAMP-mediated TLR4 activation may also be at play. C34, a small molecule TLR4 antagonist also blunted pro-inflammatory cytokine secretion in LPS-challenged BMDMs in vitro. Intraperitoneal, but not oral, administration of C34 protected mice from DSS-induced anemia with improved RBC, HB, HCT, reticulocytes in blood and erythrocyte numbers in the BM.

Consistent with this, extramedullary erythropoiesis, endotoxemia and systemic inflammation were abated after DSS in C34 intraperitoneally injected mice, suggesting that TLR4 inhibitors have therapeutic potential against IBD-associated anemia. The fact that C34 route of administration (intraperitoneal versus oral) strongly influences the protective effect against anemia could be due to different C34 biodistribution in the BM, spleen and peritoneal cavity around the gut and this warrants further investigation.

One limitation of our study is that we did not examine the contribution of DAMPs on IBD-associated anemia. TLR-4 can also be activated by endogenous DAMPs such as HMGB1, calprotectin, lactoferrin, S100 proteins, heat shock proteins, ATP, IL-1α, IL-33 and hyaluronan, which can also activate TLR4 under non-infectious conditions to induce tissue repair^59–62^ and may fuel a sterile inflammation in IBD. In line with this, HMGB1 has been suggested as a key mediator of DSS-induced colitis as antagonists targeting HMGB1 significantly alleviated DSS-induced colitis severity and inflammation.^63^ Although our study suggests endotoxemia is a key factor in IBD-associated anemia, potential contribution of DAMPs such as HMGB1 cannot be ruled out and would benefit from further investigation. However, our results in mice with conditional deletion of the *Tlr4* gene in tissue-resident macrophages as well as TLR4 antagonist administration suggest that TLR4 may play a key role in driving colitis-associated anemia.

Taken together, our work reveals new insights on IBD-associated anemia with a functional link between endotoxemia and anemia in IBD. Our work suggests that activation of TLR4 on tissue-resident Mφ drives anemia and inflammation in IBD and that TLR4 antagonists are promising therapeutic avenues for correcting IBD-associated anemia.

## Acknowledgements

This work was supported by American Society of Hematology (ASH) Global Research Award funded by ASH, Mater Research Future Leaders Fellowship funded by Mater Foundation and Leading Innovations through New Collaborations (LINC) Grant funded by the Translational Research Institute (TRI) and Mater Research Ltd. This research was carried out at TRI, Brisbane. Translational Research Institute is supported by a grant from the Australian Government. We thank TRI flow cytometry, histology, microscopy core and animal facilities. Prof Tomas Ganz (University of California, Los Angeles, USA) mentored K.B.’s ASH Global Award and critically reviewed this manuscript. Allison Pettit (Mater Research, University of Queensland, Australia) provided *Siglec1_Cre_* and Siglec1^Cre^:Zsgreen mouse model and assisted with conceptualization of this study. Yifu Tang (Sichuan Real & Best Biotech Co., Ltd. Sichuan Sheng, China) provided technical assistance for Perls’ Prussian blue iron staining.

