## Supplementary method for "Endotoxemia and TLR4 via tissue resident macrophages triggers anemia in mouse model of colitis"

**Supplemental methods**

**Mice treatment with colistin or TLR-4-IN-C34 (C34)**

Mice treated with DSS were injected intraperitoneally with 20 mg/kg colistin sulfate (Cat#C4464, SigmaAldrich) together with 400 mg/kg L-methionine^1^ (Cat#M5308, SigmaAldrich) or 1 mg/kg C34 (Cat# HY-107575, MedChem Express) once daily for the first 5 days and twice daily on day 6 and 7 of DSS treatment.

**Tissue harvest**

At the endpoint of experiments, mice were anesthetized with isofluorane and 0.5-1mL of blood collected into heparinized tubes by cardiac puncture before cervical dislocation. Spleen, bone marrow (BM), liver, duodenum and colon tissues were collected. The BM of one femur was flushed into 1 mL ice-cold PBS containing 2% newborn calf serum (NCS) and 2 mM EDTA using a 1mL syringe mounted with a 23G needle. Cells were washed in this buffer for subsequent flow cytometry analyses.

Blood samples for flow cytometry were lysed by a 6-minute incubation in 5 volumes of 10 mM NaHCO_3_, 150 mM NH_4_Cl, 1 mM EDTA pH=7.4 red cell lysis buffer at room temperature and then washed twice with PBS + 2% NCS+2 mM EDTA.

Spleens were weighed and then dissociated in 3 mL PBS 2% NCS using a GentleMACS Dissociator tissue homogenizer with matching C tubes (Miltenyi Biotec, Macquarie Park, Australia) on “spleen 3” setting, twice. In some experiments, 10-20 mg of spleen was either fixed in 4% paraformaldehyde (PFA) for Prussian blue histology staining or snap frozen on dry ice for tissue iron measurement.

Duodenum and liver were snap frozen for RNA or tissue iron measurement. Colon length and weight were measured, and colons were washed once in PBS. For Hematoxylin and Eosin histology, colon rolls were fixed in 4 % PFA. For cytokine secretion measurement, ~0.7-1 cm pieces of distal colon were cut and washed in PBS and cultured in 96 well plate containing 200 µL of RPMI-1640 medium supplemented with 10 % fetal bovine serum (FBS), penicillin and glutamine and incubated overnight in a 37ºC, 5 % CO_2_ incubator.^1^ The next morning, supernatants were collected and centrifuged at 20,000 g, 10 minutes, 4ºC and stored at -80ºC until analysis.

For plasma collection, blood samples were centrifuged twice (serially) at 800g for 10 mins and the double clarified plasma collected, aliquoted and stored at -80°C until analysis. For serum, blood was collected by cardiac puncture into tubes without anticoagulant, left to clot for 30 minutes at RT, centrifuged twice (serially) at 800 x g for 10 minutes at 4 °Cand serum collected, aliquoted and stored at -80ºC until analysis.

Blood, spleen and BM samples were counted on Mindray BC-5000 Vet Auto hematology analyzer (Biomedical Electronics Co. LTD., China).

**Isolation of peritoneal cells**

Following blood collection mice were euthanized by cervical dislocation and sterilized with 80 % ethanol. A small incision was made over the bottom half of the abdomen with scissors and underlying abdominal wall was exposed by retraction. Ten mL of sterile PBS was injected into the caudal half of the peritoneal cavity using a 25-gauge needle (beveled side up). Peritoneal cells were collected by slowly withdrawing PBS. Cells were centrifuged at 370g, 4ºC for 5 min and resuspended in 1 mL PBS + 2% NCS + 2 mM EDTA.

**Red blood cell half-life measurements**

In vivo RBC half-life span was measured by in vivo RBC surface biotinylation adapted from previously published protocol.^2^ Briefly EZ-Link Sulfo-NHS Biotin (ThermoFisher Scientific) was dissolved in PBS at a concentration of 1 mg biotin per 300 µL injection. Mice were then injected with the biotinylation reagent by retro-orbital intravenous injection under anesthesia 1-hour prior to DSS administration. About 10-20 µL of whole blood was collected at 24 h (day 0) and 72 h (day 3) from the tail vein and by cardiac puncture at the end point at 168 h (day 7) post biotinylation. Cell surface biotin was detected by flow cytometry after staining of whole blood with streptavidin-APC and Ter119-FITC antibody.

**Flow cytometry analysis**

All fluorescent antibodies used in this study are listed in supplemental Table 1. Erythroid populations in BM and spleen were measured by flow cytometry by staining samples in mouse CD16/CD32 hybridoma 2.4G2 supernatant containing fluorescent antibodies for markers for Ter119, CD44 and CD45, together with viable fluorescent DNA intercalant Hoechst33342 (Hoe; Sigma-Aldrich) as previously described.^3^ Fixable viability stain (FVS)700 (BD Bioscience) was added to all stained samples for dead cell exclusion. ^3^ Cells were washed once in PBS + 2% NCS + 2 mM EDTA.

Blood reticulocytes were stained using thiazole orange dye as previously described^5^ with a few modifications. In brief, a stock solution of thiazole orange (1mg/mL) was prepared in methanol. A working solution of 1:10,000 was prepared in PBS containing 0.02% NaN_3_ and 2 mM EDTA. 1.25 µL of whole blood was stained with 250 µL thiazole orange working solution and PercPCy5.5 anti-mouse Ter119 antibody and stained at RT for 1 h. Thiazole orange fluorescence was detected in the FITC channel.

To measure TLR4 expression, lysed blood and peritoneal cells were stained in mouse CD16/32 hybridoma 2.4G2 supernatant containing fluorescent antibodies for mouse CSF1R, CD11b, CD3ε, B220, Ly6C, Ly6G and TLR4 for 40 min on ice. For measuring the effect of DSS on the inflammatory activation state of different monocytes, blood, BM and spleen cells were stained in mouse CD16/32 hybridoma 2.4G2 supernatant containing fluorescent antibodies for mouse CD11b, CSF1R, CD64, CD86, I-A/I-E, CCR2, CX3CR1, LY6C and LY6G for 40 min on ice. Antibodies detailed in Supplementary Table 2. Samples were analyzed on a Cytoflex (Beckman Coulter) flow cytometer equipped with 640nm, 561nm, 488nm, and 405nm lasers.

For BrdU staining, bone marrow and spleen cells were surface stained in mouse CD16/CD32 hybridoma 2.4G2 supernatant containing fluorescent antibodies for Ter119, CD44 and CD45. Cells were washed, fixed, permeabilized and incubated with DNAse using BrdU-kit (Cat # 556028, Water BioSciences) as per manufacturer instructions. Following incubation with DNAse and washing, cells were stained with anti-BrdU antibody and 7-aminoactinomycin D (7-AAD). BrdU incorporation into different cell cycle phases (G0/G1, S or G2/M) was analyzed by 7-AAD intensities by flow cytometry as previously described^6^ and by following manufacturer instructions (Water BioSciences).

Uncompensated FCS files were analyzed using FlowJo 10.9 and 10.10 software following compensation with single color antibody stains (Tree Star, Ashland, OR) following post-hoc compensation with single color controls.

**In vitro treatment of bone marrow derived macrophage (BMDM) with colistin sulfate and TLR4-IN-C34**

BMDMs were prepared by flushing the BM of 1 femur with 1 mL of PBS+2 % NCS +2 mM EDTA. Cells were transferred to 50 mL tubes containing 9 mL PBS+2% NCS + 2 mM EDTA and centrifuged at 370 x g, 4ºC for 5 minutes. Cells were resuspended in RPMI-1640 medium supplemented with 10% FBS, 100ng/mL recombinant human macrophage colony-stimulating factor (CSF)-1 kindly donated by Prof David Hume (Mater Research-UQ), penicillin and glutamine. Cells were seeded in one 10 cm diameter Sterilin™ dishes (ThermoFisher Scientific, cat#SLN109) and cultured at 37ºC with 5% CO_2_. After 4 days, cultures were topped up with 4 ml fresh complete medium. At day 7, BMDMs were harvested on ice with PBS containing 4 mM EDTA. One hundred thousand cells/well were seeded in 96 well plate and incubated overnight at 37ºC with 5% CO_2_. The next morning, cells were pre-incubated with 25 μM C34 (Cat # HY-107575, Med Chem Express; stock prepared in DMSO) or an equivalent volume of DMSO for 30 min and then with LPS (Cat # L4391, Sigma-Aldrich; 100 ng/mL in PBS) for 4 h. Colistin sulfate (Cat #C4461, Sigma-Aldrich; 2 µg/mL in PBS) was preincubated with LPS for 30 minutes before being added to cells and cells were incubated for 4 hr. Supernatants were collected and stored at -80 ºC until analysis.

**Cytokine profile**

Human cytokine concentrations were measured in archived non-IBD control, CD and UC diluted serum samples (1:2) using LEGENDplex^TM^ inflammation panel 1 (13-lex; Biolegend, Cat#740808) following manufacturer’s instructions. Mouse cytokine concentrations were measured in diluted plasma (1:2), BMDM and colon supernatants using LEGENDplex^TM^ mouse inflammation panel (13 plex; BioLegend, Cat # 740446) following manufacturer’s instructions. Cytokine data were acquired on a CytoFLEX flow cytometer. Cytokine concentrations were analyzed using BioLegend’s LEGENDplex™ data analysis software (BioLegend).

**Prussian blue staining and quantification**

Ferric iron (Fe^3+^) stored in spleen and liver was detected using a Prussian blue staining method. A working solution of Prussian blue was prepared by mixing equal volumes of 20 % hydrochloric acid and 10% potassium ferrocyanide (Cat # P3289, Sigma-Aldrich) solution in water immediately before use. Paraffin-embedded tissue blocks were sectioned at 5μm thickness, rehydrated in xylene and sequentially in 100%, 95% and 70 % ethanol. Tissues were then stained with Prussian blue working solution for 20 minutes. Subsequently slides were washed in distilled water, counterstained with 0.1 % Nuclear Fast Red (provided by TRI Histology Core Facility) for 5-7 minutes, followed by rinsing in distilled water for 10 minutes. Slides were dehydrated sequentially in 95% and 100% ethanol, then in xylene, with two changes each. Slides were mounted with Cytoseal™ 60 mounting medium (ProSciTech). Images were captured on Evident VS120 slide scanners (Olympus) with 40X objective lens under bright field. Images were quantified in QuPath version 0.5 and number of positive Prussian Blue per filed was calculated.

**Tissue non-heme iron measurements**

Tissue non-heme ferric iron (Fe^3+^) was measured using a bathophenanthroline-based colorimetric assay as previously described.^6^ Briefly, wet weight of liver or spleen tissues were recorded followed by tissue drying at 65°C for 48 hr. Dried weight of each sample was recorded. Dried samples were then digested at 65ºC for 20 hr in the acid mixture containing 10g of trichloroacetic acid dissolved in 37% hydrochloric acid. After cooling, 500 µL of the clear supernatant (acid extract) was transferred into fresh Eppendorf tubes and immediately used for the colorimetric assay. Saturated sodium acetate stock, chromogen reagent containing 4,7-dipheny-1,10-phenanathroline-disulfonic acid disodium salt trihydrate chromogen (Cat#11890, Sigma-Aldrich) and thioglycolic acid, iron standard and blank were prepared as previously described.^6^ In brief, working chromogen reagent (WCR) was prepared by adding one volume of chromogen reagent to 5 volumes of saturated sodium acetate and 5 volumes of deionized water (diH_2_O). Samples (45 µL), acid blank (diH_2_O); 45 µL) and iron standard (22.5 µL standard + 22.5 µL diH_2_O) and 150 µL of WCR were added to a flat-bottom 96 well plate. Chromogen reaction was prepared by incubating plate for 15 min at RT. Absorbance was measured at 535 nm using a spectrophotometer (Thermo Fisher Scientific), and non-heme tissue iron was calculated with the following equations.

$$Tissue iron \left( \mu g/g dry tissue \right)=\frac{AT-AB}{AS-AB} X\frac{Fes}{W} x\frac{\frac{Vrv}{Vsmp}xVf}{Vrv/Vstd}$$

AT = absorbance of test sample

AB = absorbance of acid blank

AS = absorbance of standard

Fes = iron concentration of working iron standard solution (µg Fe/mL)

W = weight of dry tissue (g)

Vsmp = sample volume (mL)

Vf = final volume of acid mixture after overnight incubation at 65ºC (mL)

Vstd = iron standard voume (mL)

Vrv = final reaction volume (mL)

**Serum iron assay**

Iron concentration, total iron-binding capacity (TIBC), unsaturated iron-binding capacity (UIBC) were measured in mouse serum using the Pointe Iron/TIBC Reagent kit (Pointe Scientific, #23-666-320) according to the manufacturer’s instructions. For serum iron, 25 µL of undiluted serum and 125 µL of iron binding reagent was added to flat-bottom 96 well plates. For UIBC, 25 µL of undiluted serum, 25 µL of iron standard (500 µg/dL) and 100 µL of UIBC buffer reagent was added to 96 well plates. Absorbance (A; A1 reading) was measured at 560 nm using a spectrophotometer (Thermo Fisher Scientific). Then 2.5 µL of iron colour reagent was added to each well and incubated for 30 min at 37ºC and absorbance was measured at 560 nM (A2 reading). Serum iron, UIBC, TIBC and % transferrin saturation were calculated using formula below.

$$Serum iron \left( \mu g/dl \right)= \frac{A2Test-A1Test}{A2STD-A1STD} x conc of STD$$

$$UIBC \left( \mu g/dl \right)=Conc of STD-\left( \frac{A2Test-A1Test}{A2STD-A1STD} \right)x conc of STD$$

$$TIBC (\mu g/dl)=Serum iron+UIBC$$

$$\% Transferrin saturation=\left( \frac{serum iron}{TIBC} \right)x 100\%$$

A= absorbance, STD = standard, Test = sample

**Hematoxylin and Eosin (H&E) staining and quantification**

For histology scoring, PFA fixed and paraffin embedded colon rolls were sectioned at 5 µm thickness and stained with H&E by the TRI Histology Core facility. Blind assessment of histologic inflammation was performed as previously described.^1^ Briefly, distal colons were scored (score = 0 for no severity and ≥5 for severely damaged colons) for increased leukocyte infiltration, neutrophil counts, depletion of goblet cells, crypt abscesses, aberrant crypt architecture, increased crypt length and epithelial damage and ulceration.

**Immunofluorescent staining and imaging**

Paraformaldehyde fixed colon rolls were embedded in Tissue-Tek O.C.T. and were kept frozen at -80ºC. Frozen sections were cut at 5µm thickness using a Leica CM1950 cryostat (Leica Biosystems). Slides were air dried at RT for 1 hour, then rehydrated in 0.1% Triton X‑100 in Tris‑buffered saline (TBS) for 5-10 min prior to staining. To block non-specific binding, sections were blocked for 1 hr at RT using a blocking solution consisting of 3 % BSA, 10% normal goat serum, 5 % normal donkey serum and 0.1 % Triton X-100 in TBS (TBST), then washed by dipping in TBS. Following blocking, sections were incubated for 1.5 h at RT with a rabbit anti‑IBA‑1 (Fujifilm Wako Pure Chemical Corporation) and rat anti‑mouse CD169 (BioLegend) primary antibodies diluted in TBST, followed by washing in TBS. Slides were then incubated for 40 min at RT in the dark with donkey anti‑rabbit IgG Alexa Fluor 555 (Thermo Fisher Scientific) and donkey anti‑rat IgG Alexa Fluor 647 (Thermo Fisher). After washing in TBS, sections were mounted in Fluoroshield^TM^ with DAPI histology mounting medium (Sigma‑Aldrich, #F6057). Full antibody details are provided in the supplemental Table 1. Images were captured on Evident VS200 slide scanner (Olympus) with 20 X objective lens. Images were uploaded in QuPath version 0.5 and TIFF files were exported.

**Statistical analysis**

Statistical analyses were calculated using GraphPad Prism (v10.4). Data are presented as mean ± SD unless specified otherwise. Statistical tests and p values are specified in the figure legends or figures. P value <0.05 was considered statistically significant.
