## Supplementary Table for "Endotoxemia and TLR4 via tissue resident macrophages triggers anemia in mouse model of colitis"

Supplemental Table 1. Antibodies and reagents used for flow cytometry and immunofluorescent staining

| **Reagents and resources** | **Suppliers** | **Catalog numbers** | **Clone** | **Dilution** |
| --- | --- | --- | --- | --- |
| Anti-mouse Ter119- Percpcy5.5 | BioLegend | 116228 | TER-119 | 1/200 |
| Anti-mouse Ter119-FITC | BioLegend | 116206 | [TER-119](https://www.biolegend.com/nl-be/search-results?Clone=30-F11) | 1/200 |
| Anti-mouse CD44 – APC | Biolegend | 103012 | IM7 | 1/300 |
| Anti-mouse CD45 – APCCY7 | Biolegend | 103116 | 30-F11 | 1/200 |
| Anti-mouse CSF1R (CD115)-PECY7 | BioLegend | 135524 | AFS98 | 1/200 |
| Anti-mouse/human CD11b-BV510 | BioLegend | 101245 | M1/70 | 1/200 |
| Anti-mouse/human CD11b-FITC | BioLegend | 101206 | M1/70 | 1/200 |
| Anti-mouse TLR-4/MD2 complex-PE | BioLegend | 117605 | MTS510 | 1/75 |
| Anti-mouse Ly6C-FITC | BioLegend | 128006 | HK1.4 | 1/200 |
| Anti-mouse CD3ε -Percpcy5.5 | BioLegend | 100328 | 145-2C11 | 1/200 |
| Anti-mouse CD3e -Pacific blue (PB) | BioLegend | 100333 | 145-2C11 | 1/200 |
| Anti-mouse/human CD45R (B220)-APCCY7 | BioLegend | 03224 | [RA3-6B2](https://www.biolegend.com/nl-be/search-results?Clone=RA3-6B2) | 1/200 |
| Anti-mouse LY6G-BV650 | BioLegend | 127641 | 1A8 | 1/200 |
| Anti-mouse LY6G-APCCY7 | BioLegend | 127624 | 1A8 | 1/200 |
| Anti-mouse F4/80-BV421 | BioLegend | 123131 | BM8 | 1/200 |
| Anti-mouse CD25-PECY7 | BioLegend | 101916 | 3C7 | 1/100 |
| Anti-mouse CD4-FITC | BioLegend | 100405 | GK1.5 | 1/200 |
| Anti-mouse CD62L-PE | BioLegend | 161203 | W18021D | 1/200 |
| Anti-mouse LY6C-PB | BioLegend | 128014 | HK1.4 | 1/200 |
| Anti-mouse CD16/32-Percpcy5.5 | BioLegend | 101324 | 93 | 1/100 |
| Anti-mouse CD64 (FcγRI)-APC | BioLegend | 161005 | S18017D | 1/200 |
| Anti-mouse CD86-PE | BioLegend | 105105 | PO3 | 1/200 |
| Anti-mouse CD192 (CCR2)-APC | BioLegend | 150627 | SA203G11 | 1/200 |
| Anti-mouse CX3CR1-PE | BioLegend | 149005 | SA011F11 | 1/300 |
| Anti-mouse I-A/I-E-BV510 | BioLegend | 107635 | M5/114.15.2 | 1/200 |
| Anti-BrdU-PE | BioLegend | 339812 | Bu20a | 1/20 |
| Rabbit anti-Iba1 | Fujifilm Wako Pure Chemical Corporation | 019‑19741 |  | 1/200 |
| Purified anti-mouse CD169 (Siglec-1) | Biolegend | 142402 | 3D6.112 | 1/300 |
| Donkey anti-rabbit IgG (H+L) Alexa Fluor 555 | ThermoFisher Scientific | A31572 |  | 1/600 |
| Donkey anti-rat IgG (H+L) Alexa Fluor Plus 647 | ThermoFisher Scientific | A48272 |  | 1/600 |
| Rabbit IgG isotype control | ThermoFisher Scientific | 312325 |  | 1/4000 |
| Purified Rat IgG2a | BioLegend | 400502 |  | 1/150 |
| Streptavidin-APC | BioLegend | 405207 |  | 1/200 |
| FC block (neutralizing mouse CD16/32) | Dr Louise Purton |  | 2.4G2 | 50% |
| Hoechst33342 | ThermoFisher Scientific | 62249 |  | 1/800 |
| FVS700 | BD Biosciences | 564997 |  | 1/10,000 |
| 7-AAD | ThermoFisher Scientific | A1310 |  | 1/10 |
