## Supplementary Figures for "Endotoxemia and TLR4 via tissue resident macrophages triggers anemia in mouse model of colitis"

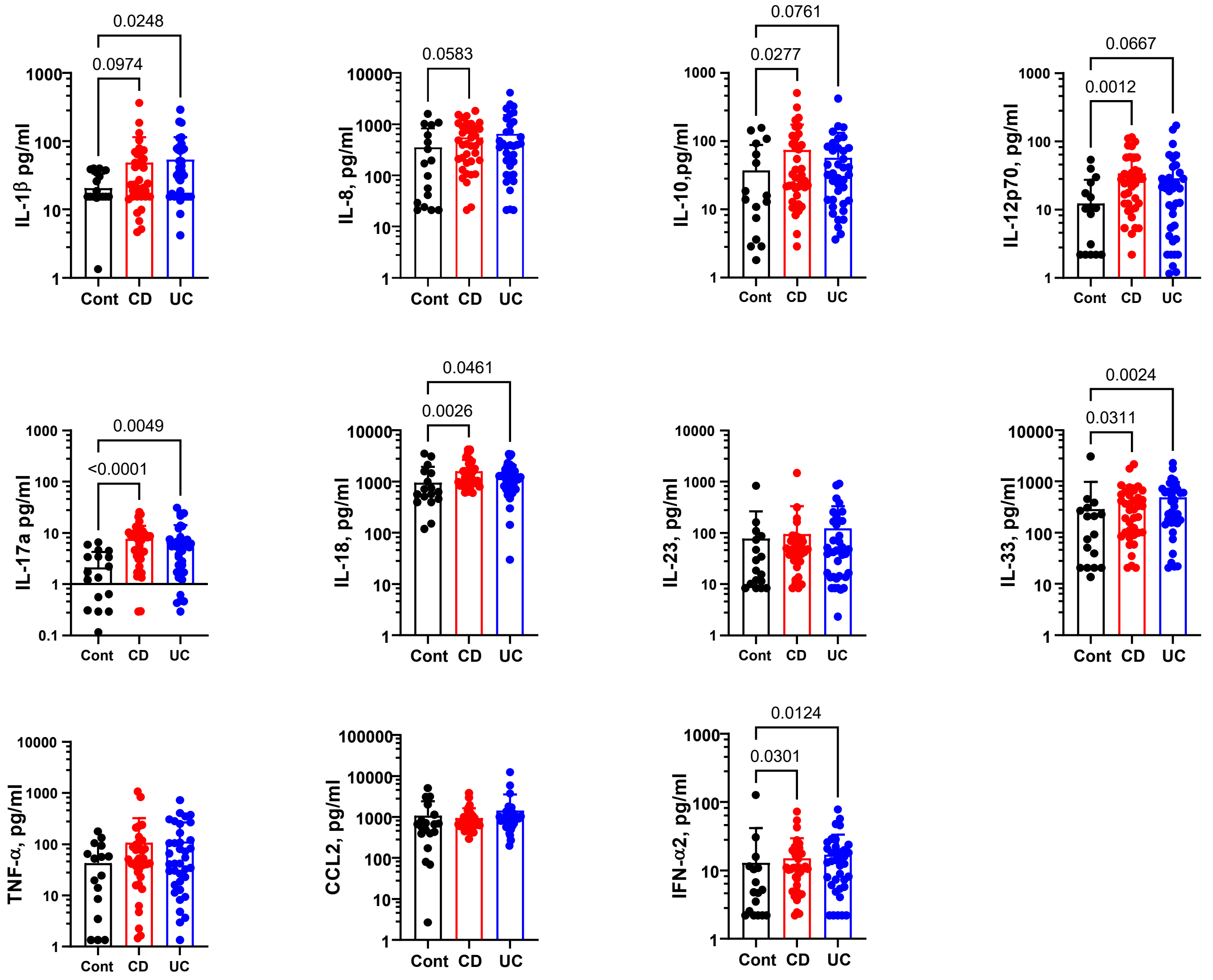


**Supplemental Figure 1. Cytokine profile of human non-IBD control (cont), Crohn's disease (CD) and ulcerative colitis (UC) patients.** Cytokine concentrations were measured in retrieved patients' serum samples by Legendplex human inflammation panel (13 plex). Each dot represents a different patient. Statistical analyses were performed by Kruskal-Wallis test followed by Dunn's post-hoc multiple comparison. Bars represent mean ± SD (n=19/cont, n=38/CD and UC group).

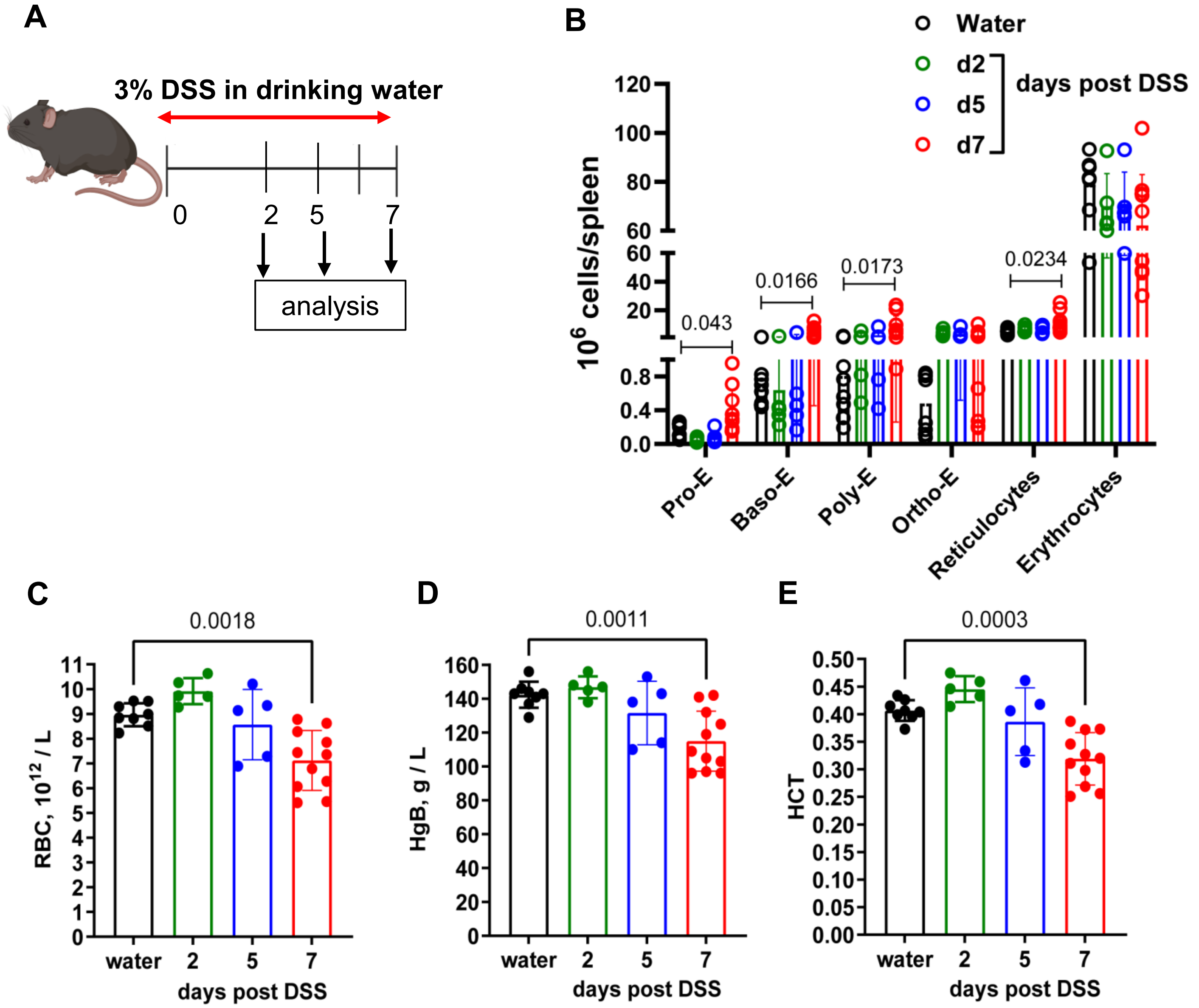


**Supplemental Figure 2. DSS-induced colitis in mice causes anemia in a time-dependent manner.** A) Schematic of 3 % DSS time course experiment. Blood and spleen were harvested at day 2, 5 and 7 post-DSS treatment. B) Number of proerythroblasts (pro-E), basophilic erythroblasts (Baso-E), polychromatic erythroblasts (Poly-E), orthochromatic erythroblasts (Ortho-E), reticulocytes and erythrocytes in the spleen were quantified by flow cytometry. C) RBC counts, D) hemoglobin (HgB) concentrations and E) hematocrit (HCT) were measured by Mindray hematology analyzer. Data are from 1 experiment (n=8/water, n=5/d2 and d5, n=10/d7 groups). Each dot is a separate mouse. Statistical significances analyses were performed by One-Way ANOVA with Sidak's multiple comparison test. Bars represent mean ± SD.

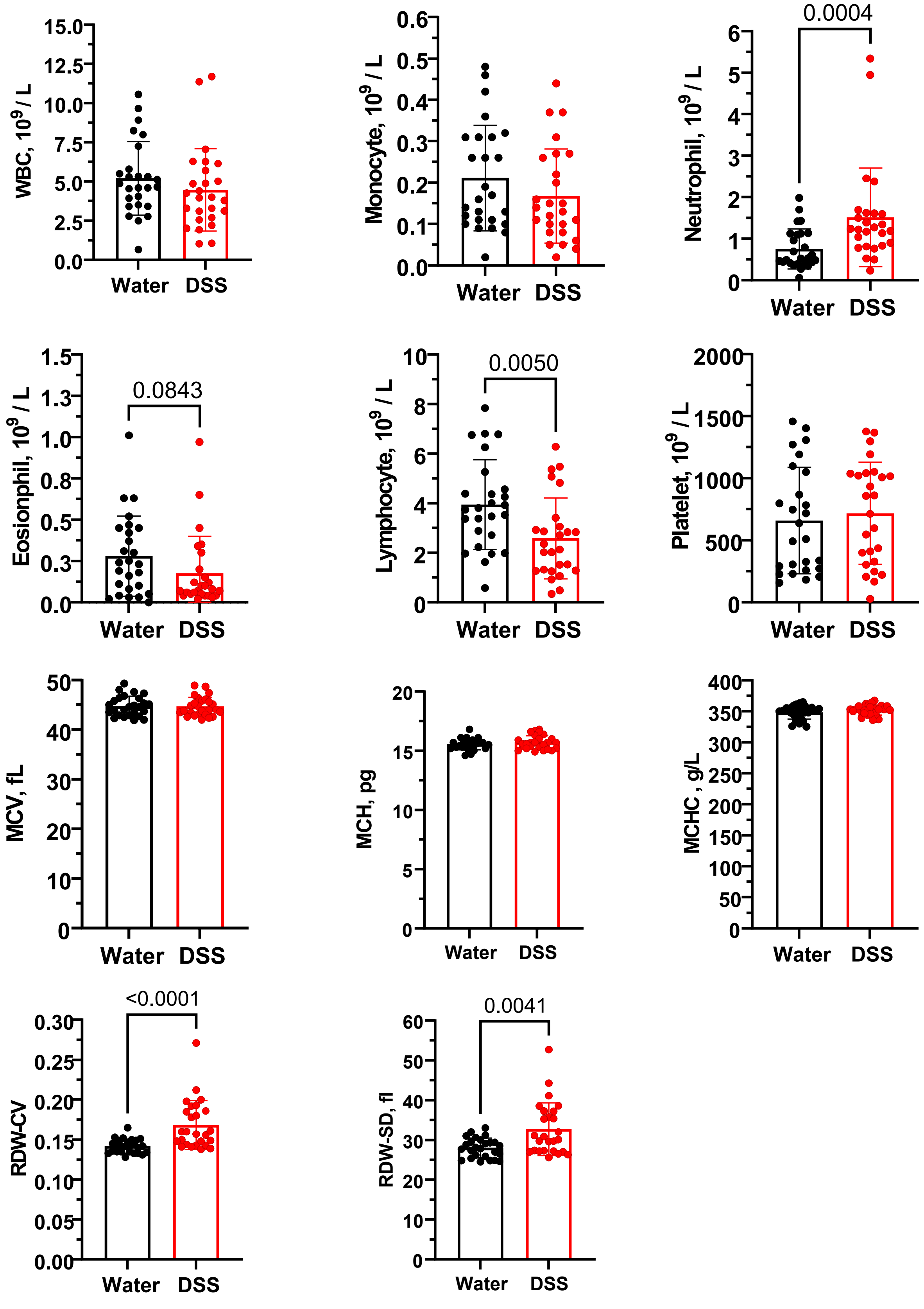


**Supplemental Figure 3. Effect of DSS-induced colitis on peripheral blood counts.** The number of white blood cells (WBC), monocytes, neutrophils, eosinophils, lymphocytes, platelet, mean corpuscular volume (MCV), mean corpuscular hemoglobin (MCH), mean corpuscular hemoglobin concentration (MCHC), red blood cell distribution width-coefficient variation (RDW-CV) and width-standard deviation (RDW-SD) were measured with a Mindray BC-5000 Vet Auto hematology analyzer. Data are pooled from 7 different experiments (n=26/group). Statistical analyses were performed by Mann–Whitney test. Each dot is a separate mouse. Bars represent mean ± SD.

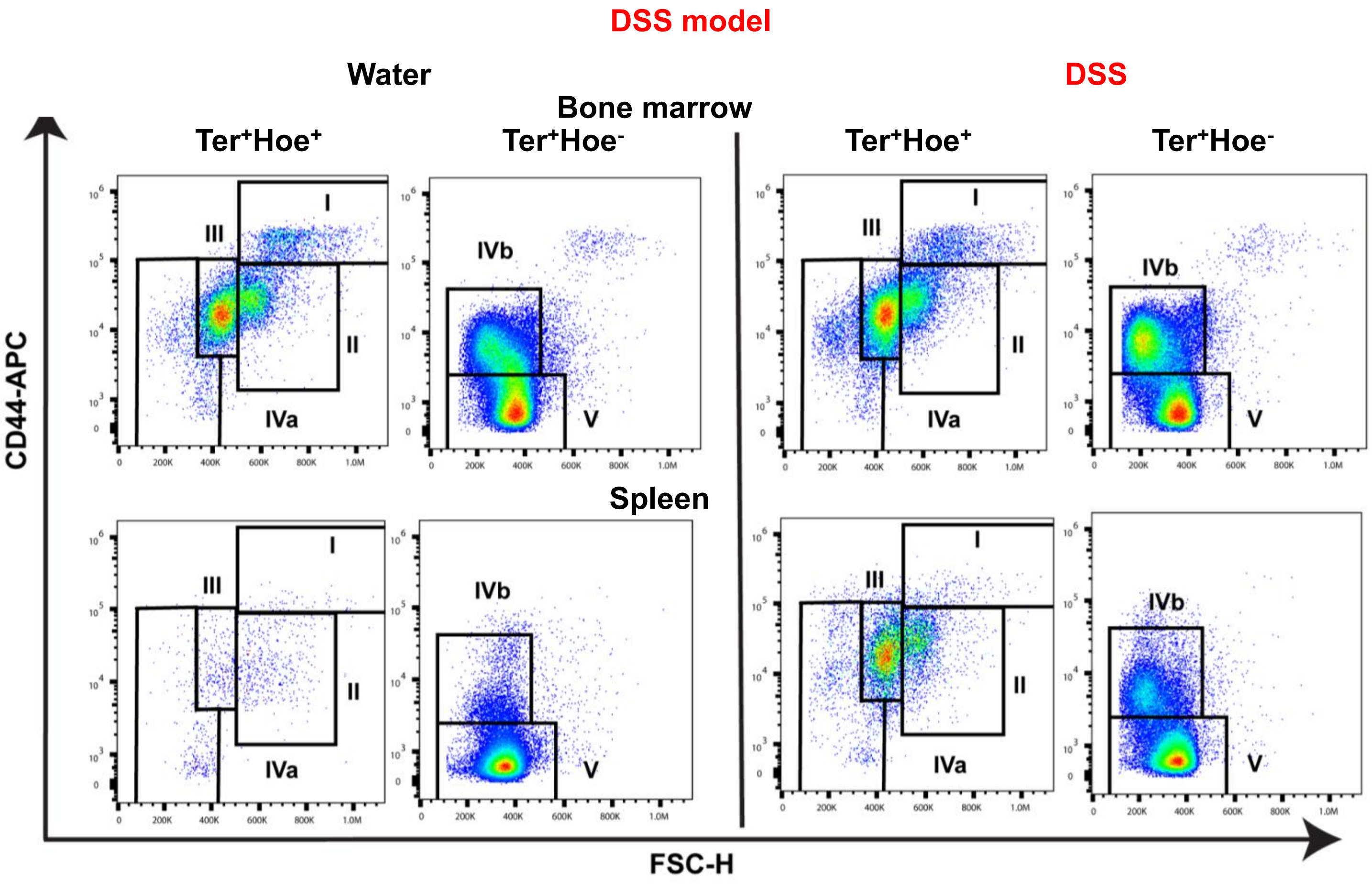


**Supplemental Figure 4. DSS-induced colitis in mouse causes anemia and increases extramedullary erythropoiesis.** Flow cytometry plots showing changes in erythropoiesis in bone marrow and spleen in response to DSS-induced colitis. This example is taken from a C57BL/6 mouse treated with plain water or 3% DSS for 7 days. Singlet cells were gated according to forward and side scatter and then gated for viable (FVS700-negative) followed by nucleated Ter119$^{+}$Hoechst (Hoe)$^{+}$ or enucleated Ter119$^{+}$Hoe$^{-}$ erythroid cells. Nucleated proerythroblasts (I; Hoe$^{+}$Ter119$^{+}$FSC$^{hgh}$D44$^{high}$), basophilic-EB (II; Hoe$^{+}$Ter119$^{+}$FSC$^{int}$CD44$^{+}$), polychromatic-EB (III; Hoe$^{+}$Ter119$^{+}$FSC$^{int}$CD44$^{low}$) and orthochromatic-EB (IVa; Hoe$^{+}$Ter119$^{+}$FSC$^{low}$CD44$^{low}$), enucleated reticulocytes (IVb; Hoe$^{-}$Ter119$^{+}$FSC$^{low}$CD44$^{low}$ ) and erythrocytes (V; Hoe$^{-}$Ter119$^{+}$FSC$^{low}$CD44$^{-}$ ).

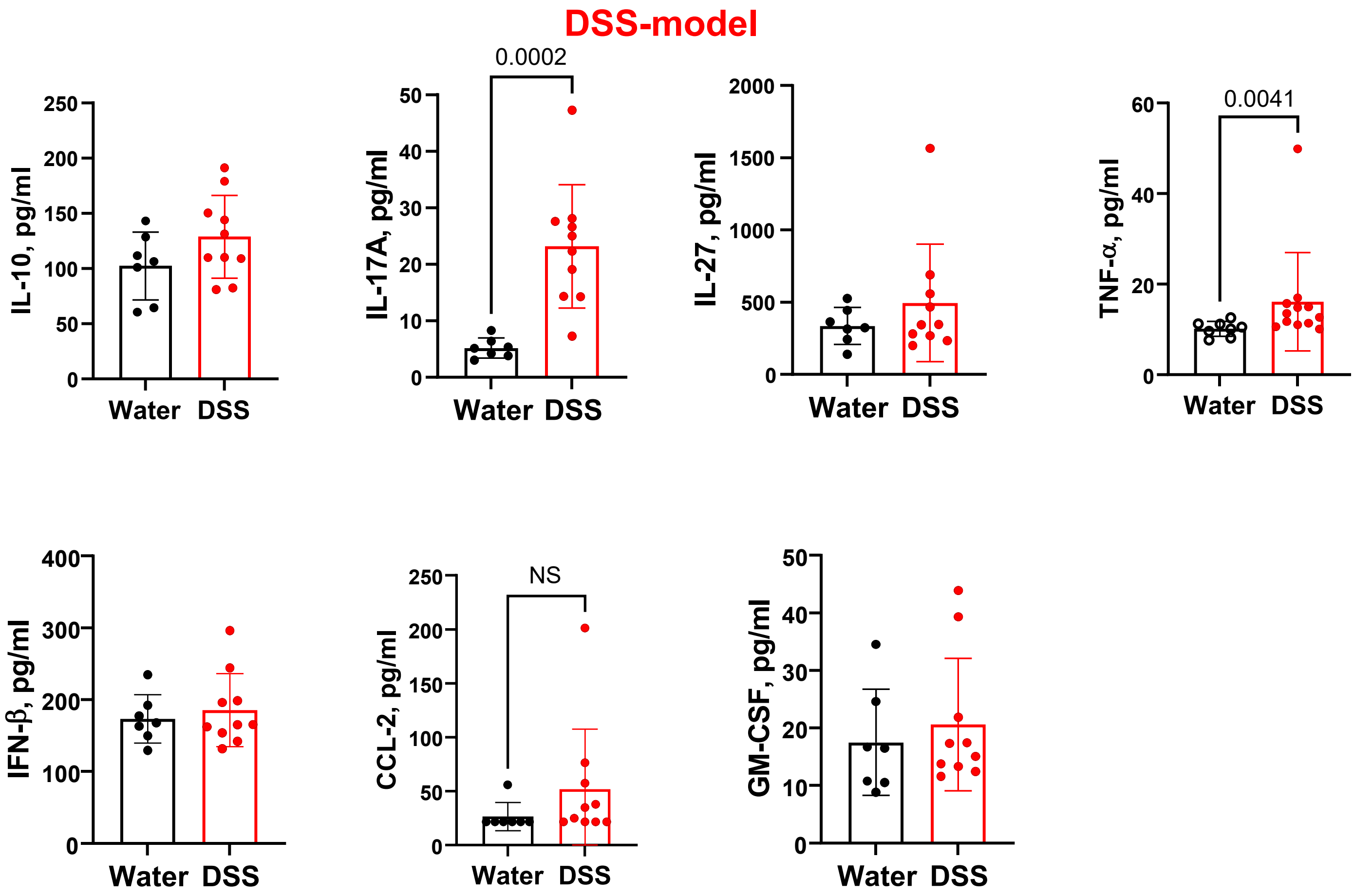


**Supplemental Figure 5. Cytokine levels in plasma of mice following DSS treatment.** Colitis was induced in mice by 3% DSS in drinking water as shown in Figure 2. Cytokine concentrations were measured in plasma samples at day 7 by Legendplex mouse inflammation panel (13 plex). Data are pooled from two different experiments. Each dot is a separate mouse. Bars represent mean ± SD (n=7 mice/water group and 10 mice/DSS group. Statistical analyses were performed by Mann-Whitney test. Bars represent mean ± SD . NS = not significant

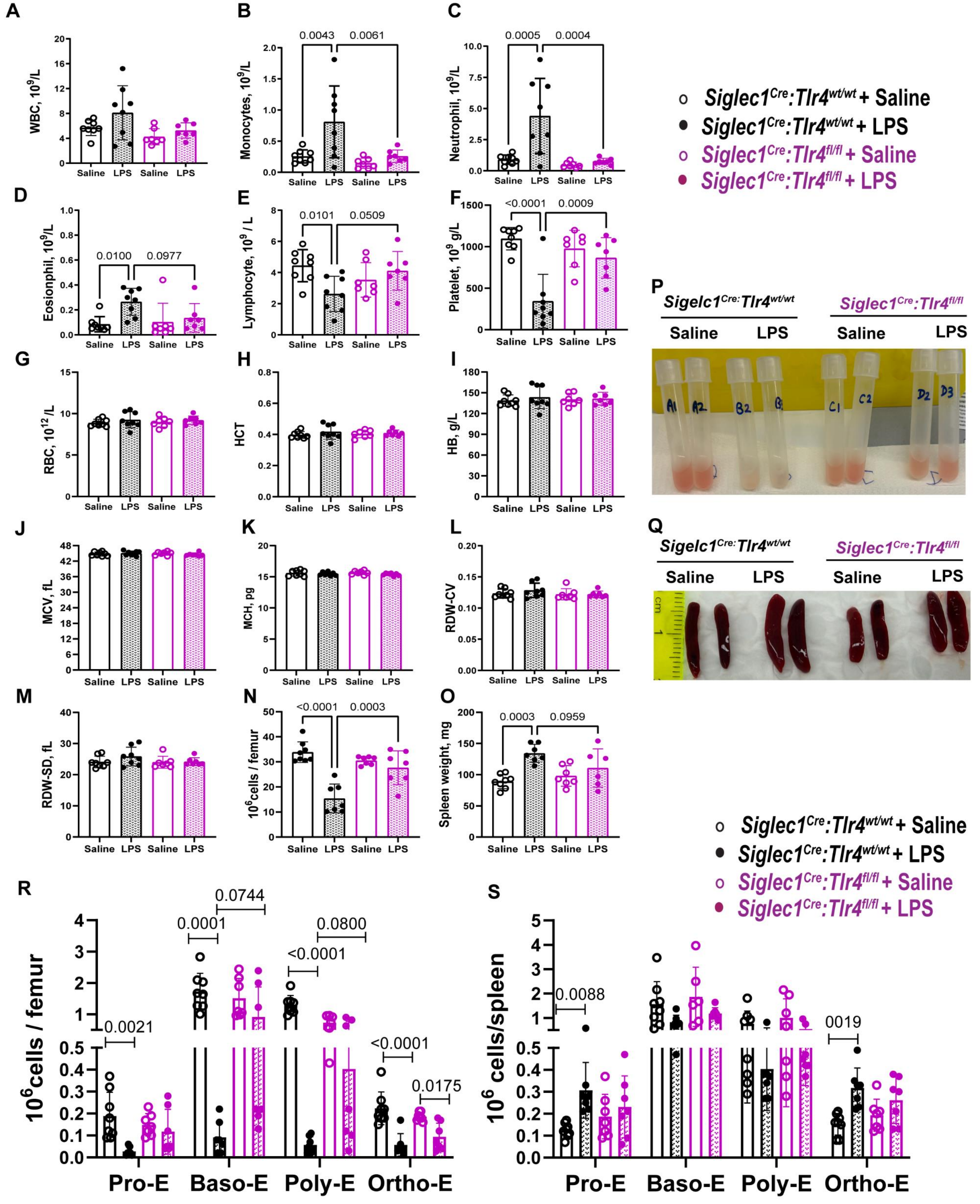

A
WBC, 10^9/L
Saline
LPS
Saline
LPS
B
Monocytes, 10^9/L
0.0043
0.0061
C
Neutrophil, 10^9/L
0.0005
0.0004
Siglec1^Cre:Tlr4^wt/wt + Saline
Siglec1^Cre:Tlr4^wt/wt + LPS
Siglec1^Cre:Tlr4^fl/fl + Saline
Siglec1^Cre:Tlr4^fl/fl + LPS
D
Eosionphil, 10^9/L
0.0100
0.0977
E
Lymphocyte, 10^9 / L
0.0101
0.0509
F
Platelet, 10^9 g/L
<0.0001
0.0009
P
Sigelc1^Cre:Tlr4^wt/wt
Siglec1^Cre:Tlr4^fl/fl
Saline
LPS
Saline
LPS
G
RBC, 10^12 /L
H
HCT
I
HB, g/L
J
MCV, fL
K
MCH, pg
L
RDW-CV
Q
Sigelc1^Cre:Tlr4^wt/wt
Siglec1^Cre:Tlr4^fl/fl
Saline
LPS
Saline
LPS
M
RDW-SD, fL
N
10^6cells / femur
<0.0001
0.0003
O
Spleen weight, mg
0.0003
0.0959
Siglec1^Cre:Tlr4^wt/wt + Saline
Siglec1^Cre:Tlr4^wt/wt + LPS
Siglec1^Cre:Tlr4^fl/fl + Saline
Siglec1^Cre:Tlr4^fl/fl + LPS
R
10^6cells / femur
0.0021
0.0001
0.0744
<0.0001
0.0800
<0.0001
0.0175
Pro-E
Baso-E
Poly-E
Ortho-E
S
10^6 cells/spleen
0.0088
0019
Pro-E
Baso-E
Poly-E
Ortho-E

**Figure 6. Conditional deletion of *Tlr4* gene in CD169$^{+}$ tissue-resident macrophages (Mφ) reduces LPS-induced changes in blood profile, improved erythropoiesis in the blood and spleen.** The number of A) white blood cells (WBC), B) monocytes, C) neutrophils, D) eosinophils, E) lymphocytes, F) platelet, G) RBC, H) HCT, I) HB, J) MCV, K) MCH, L) MCHC, M) RDW-CV) and N) RDW-SD) were measured in the blood with a Mindray BC-5000 Vet Auto hematology analyzer. N) BM cellularity was measured in the femur. O) Spleen weights. P) Photographs of mouse femoral BM flushed into 1 mL PBS and Q) spleen from saline or LPS treated *Siglec1$^{Cre}$:Tlr4$^{fl/fl}$* mice and control *Siglec1$^{Cre}$:Tlr4$^{WT/WT}$* mice. R-S) Number of proerythroblasts and erythroblasts per femur (R) or spleen (S) quantified by flow cytometry (subsets described in more detail Supplementary Figure 4). Each dot is a separate mouse. Bars represent mean ± SD (n=7-8/group) *Siglec1$^{Cre}$:Tlr4$^{fl/fl}$* mice (purple) and control *Siglec1$^{Cre}$:Tlr4$^{WT/WT}$* mice (black).

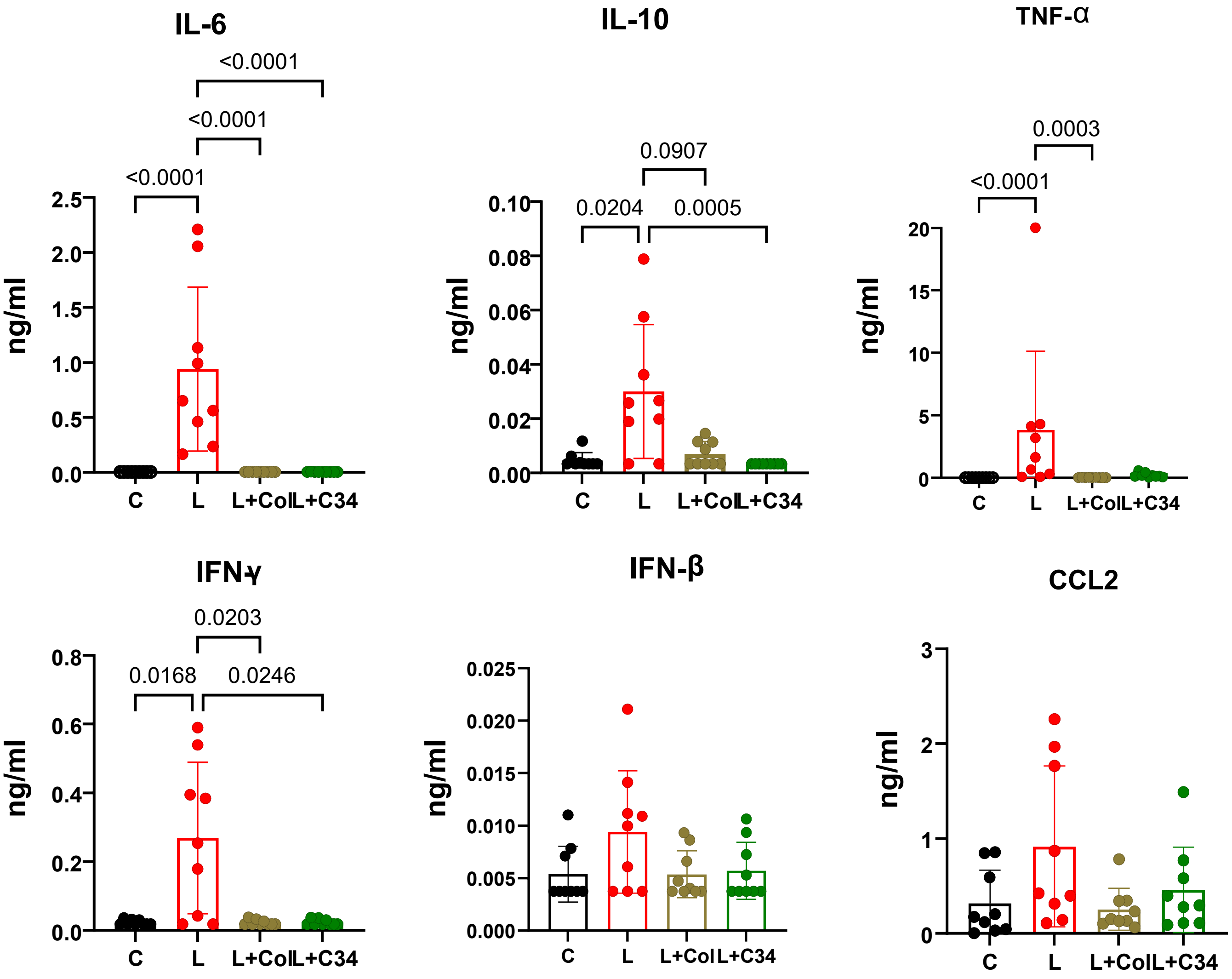


**Supplemental Figure 7. Colistin sulfate and TLR-inh-C34 block LPS-induced cytokine release by mouse bone marrow-derived monocytes (BMDM) in vitro.** BMDM (1 x $10^5$ cells/ml) were pre-incubated with 25 µM C34 for 30 min and then incubated with 100 ng/ml LPS for 4 hours. Colistin sulfate (Col) 2 µg/mL was pre-incubated with LPS at 37 ºC for 30 minutes and was then added to BMDM and incubated for 4 hours. Cytokine concentrations were measured in invitro in BMDM by Legendplex mouse inflammation panel (13 plex). Statistical analyses were performed by Kruskal-Wallis test followed by Dunn's post-hoc multiple comparison. Each dot represents a different sample. Data is pooled from 3 independent experiments. Bars represent mean ± SD (n=6-9 /group). L = LPS, L+Col = LPS + colistin sulfate and L+C34 = LPS + C34

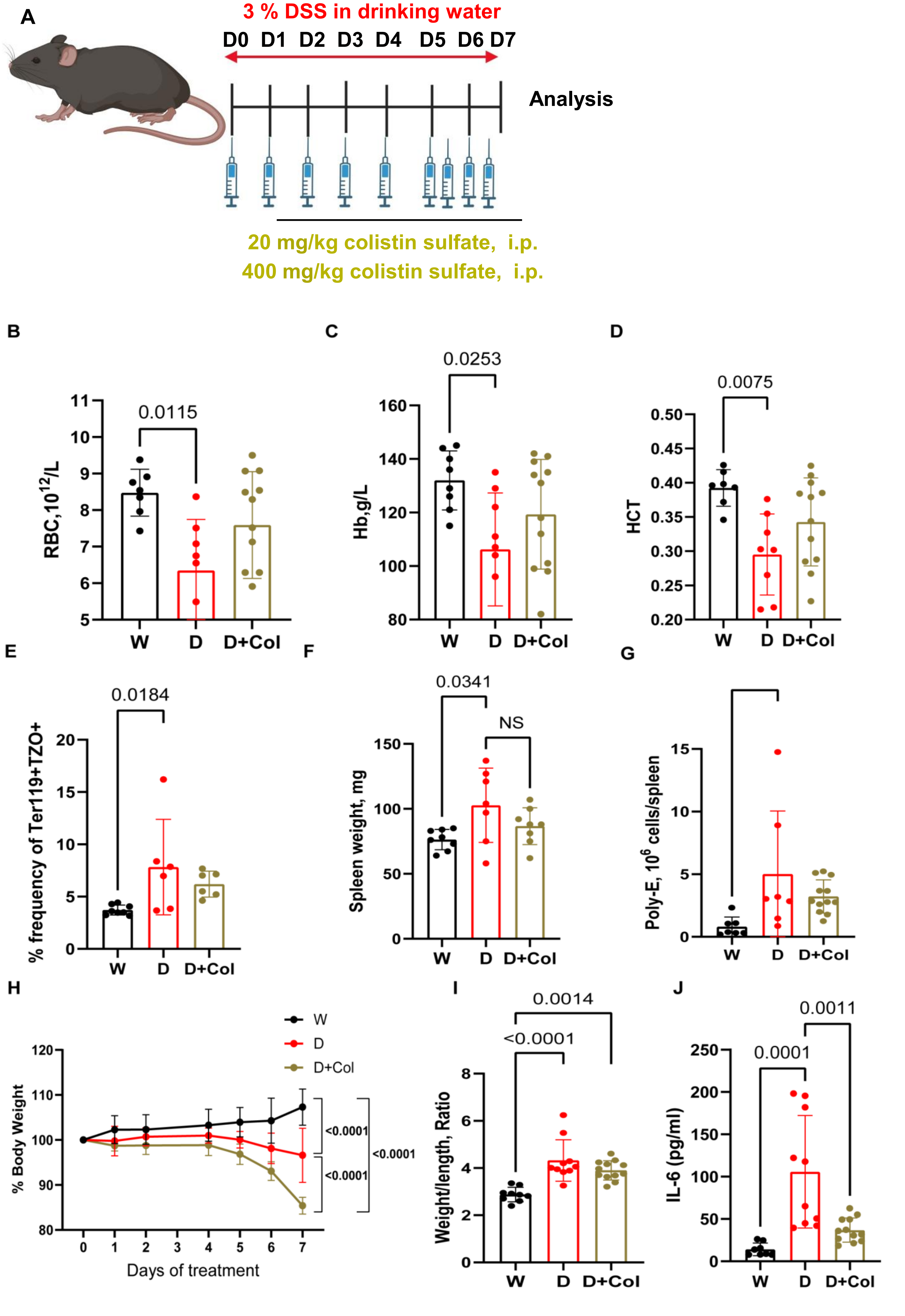
A
3 % DSS in drinking water
D0 D1 D2 D3 D4 D5 D6 D7
Analysis
20 mg/kg colistin sulfate, i.p.
400 mg/kg colistin sulfate, i.p.
B
RBC,10^12/L
0.0115
W D D+Col
C
Hb,g/L
0.0253
W D D+Col
D
HCT
0.0075
W D D+Col
E
% frequency of Ter119+TZO+
0.0184
W D D+Col
F
Spleen weight, mg
0.0341
NS
W D D+Col
G
Poly-E, 10^6 cells/spleen
W D D+Col
H
% Body Weight
Days of treatment
W
D
D+Col
<0.0001
<0.0001
<0.0001
I
Weight/length, Ratio
0.0014
<0.0001
W D D+Col
J
IL-6 (pg/ml)
0.0011
0.0001
W D D+Col

**Supplemental Figure 8. Colistin treatment does not correct DSS-induced anemia and colitis.** A) Schematic of colistin sulfate (colistin; Col), L-methionine and DSS treatments. B) RBC, C) Hb and D) HCT counts were measured in blood by Mindray hematology analyzer at day 7 of treatment with water (W), DSS (D), or DSS with Col (D+Col). E) frequency of reticulocytes ($Ter119^{+}TZO^{+}$) were quantified by flow cytometery. F) At the endpoint, spleen weights were recorded and G) numbers of polychromatic EB (Poly-E) were quantified by flow cytometery. H) Relative mouse weight loss was recorded daily. I) Colon weights and lengths were recorded and ratio were calculated for each individual mouse. J) IL-6 concentration was measured in plasma by at day7 by Legendplex. Statistical analyses were performed by One-Way ANOVA with Sidak's multiple comparison test. Data are pooled from 2 independent experiments. Each dot is a separate mouse. Bars represent mean ± SD (n = 7-9/W, 6-10/D and 6-12/ D+Col group).

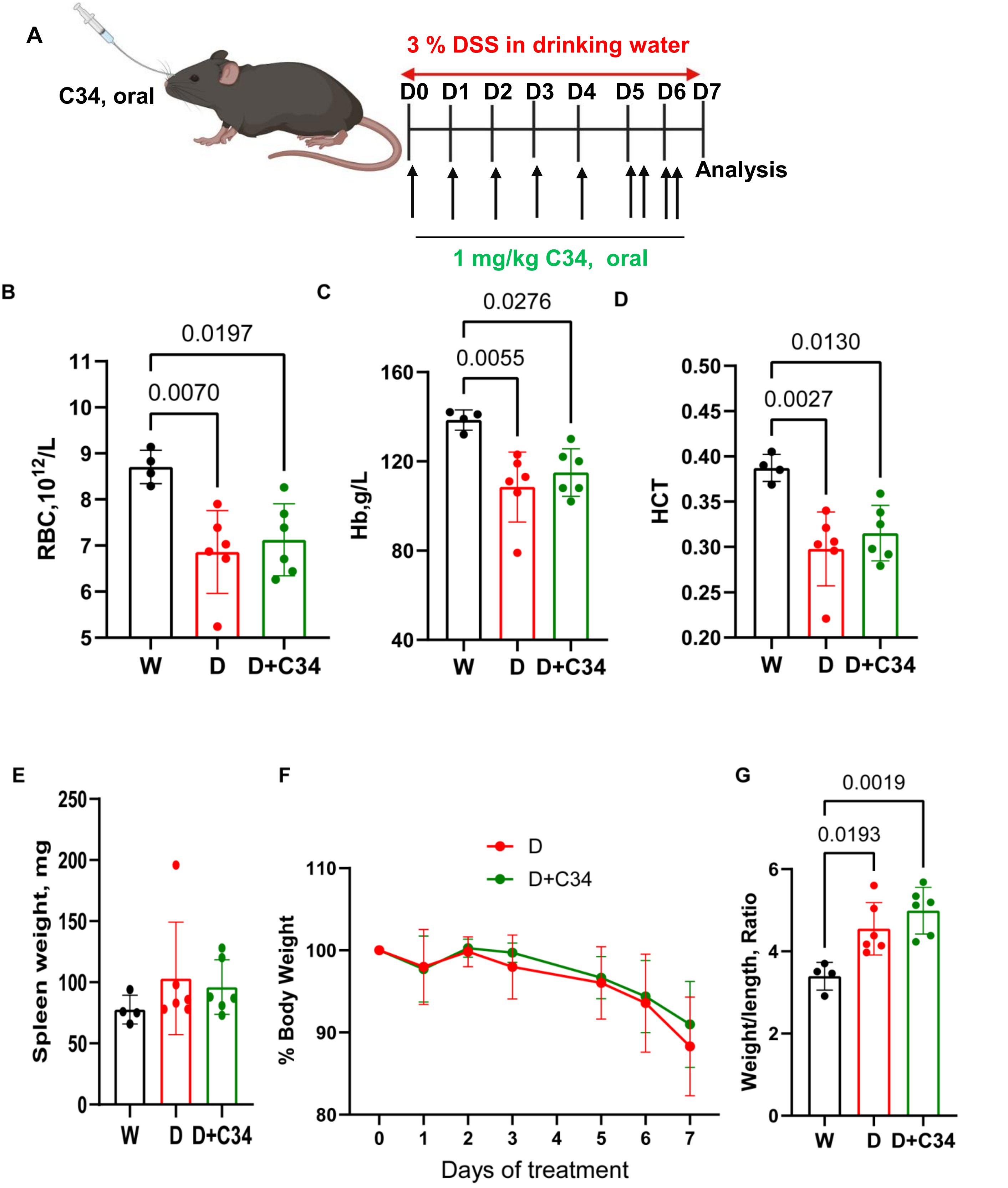


**Supplemental Figure 9. Oral C34 does not correct DSS-induced anemia and colitis.** A) Schematic of C34 and DSS treatments. Mice were given C34 via oral route for 5 days and then twice for last two days of DSS treatment. B) RBC, C) Hb and D) HCT counts were measured in blood by Mindray hematology analyzer at day 7 of treatment with water (W), DSS (D), or DSS with C34 (DSS+C34). E) Spleen weights were recorded at the endpoint. F) Relative mouse body weight loss was recorded daily. G) Colon weights and lengths were recorded and ratio were calculated for each individual mouse. Statistical analyses were performed by One-Way ANOVA with Sidak's multiple comparison test. Data are pooled from 1 experiment. Each dot is a separate mouse. Bars represent mean ± SD (n = 4/W and 6/D and D+C34 group).

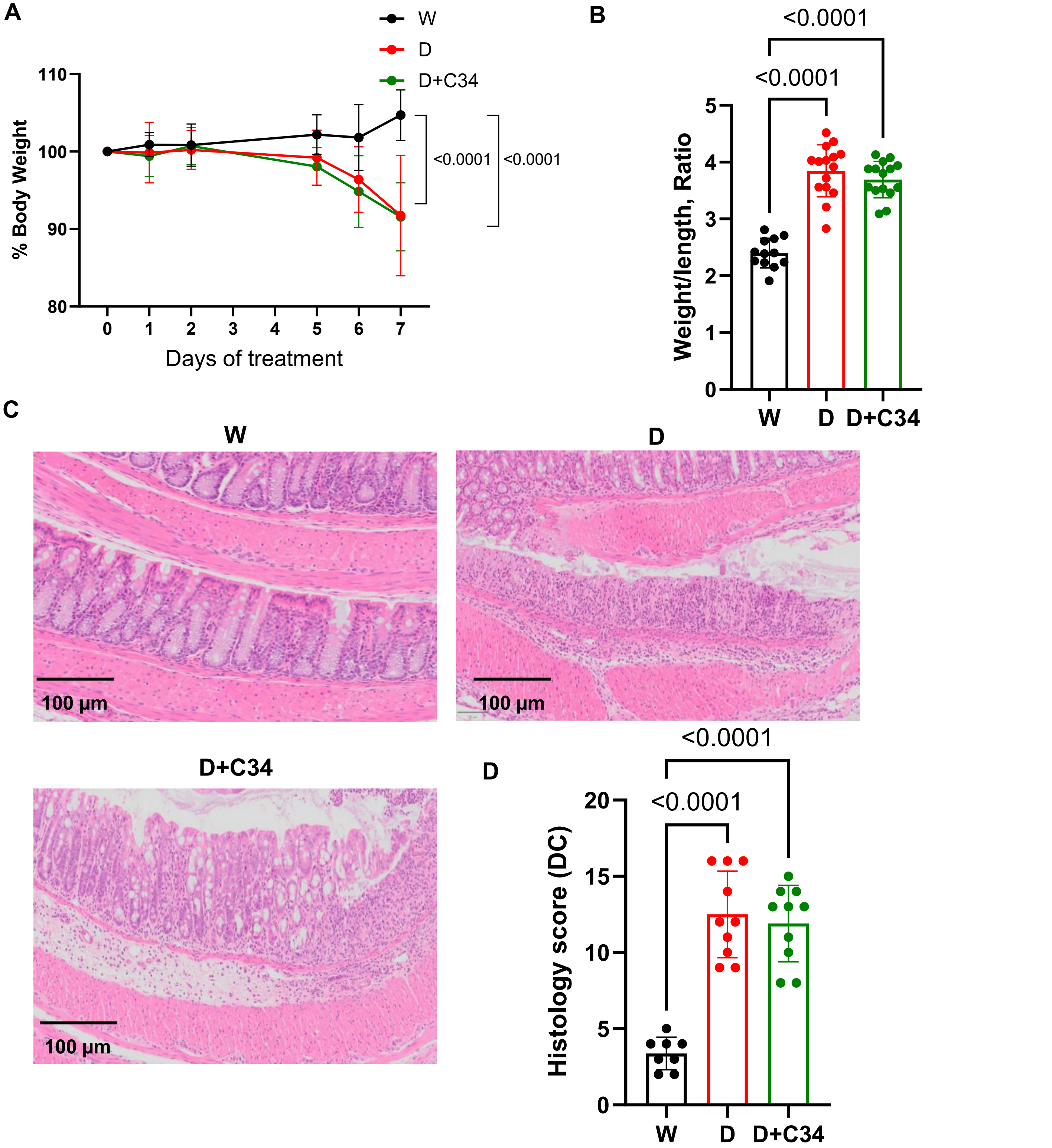


**Supplemental Figure 10. Intraperitoneal administration of C34 does not alleviate DSS-induced colitis.** Mice challenged with DSS were administered once daily with C34 via i.p. route for 5 days and then twice for last two days of DSS treatment as in Figure 7A. A) Relative mouse body weight loss was recorded daily. Statistical significances were calculated by Two-Way ANOVA with Sidak's multiple comparison test (n = 12/W, 14/D and D+C34 group). Each dot represents mean ± SD. B) Colon weights and lengths were recorded and ratio were calculated for each individual mouse. Statistical analyses were performed by One-Way ANOVA with Sidak's multiple comparison test. Data are pooled from 3 independent experiments (n = 12/W, 14/D and D+C34 group). Each dot is a separate mouse. Bars represent mean ± SD. C) Representative hematoxylin and eosin staining of mouse colon histology sections. D) Quantification of histology scores of distal colon (DC). Statistical analyses were performed by One-Way ANOVA with Sidak's multiple comparison test. Data are pooled from 2 independent experiments n = 8/W, 10 /D and D+C34 groups). Each dot is a separate mouse. Bars represent mean ± SD.

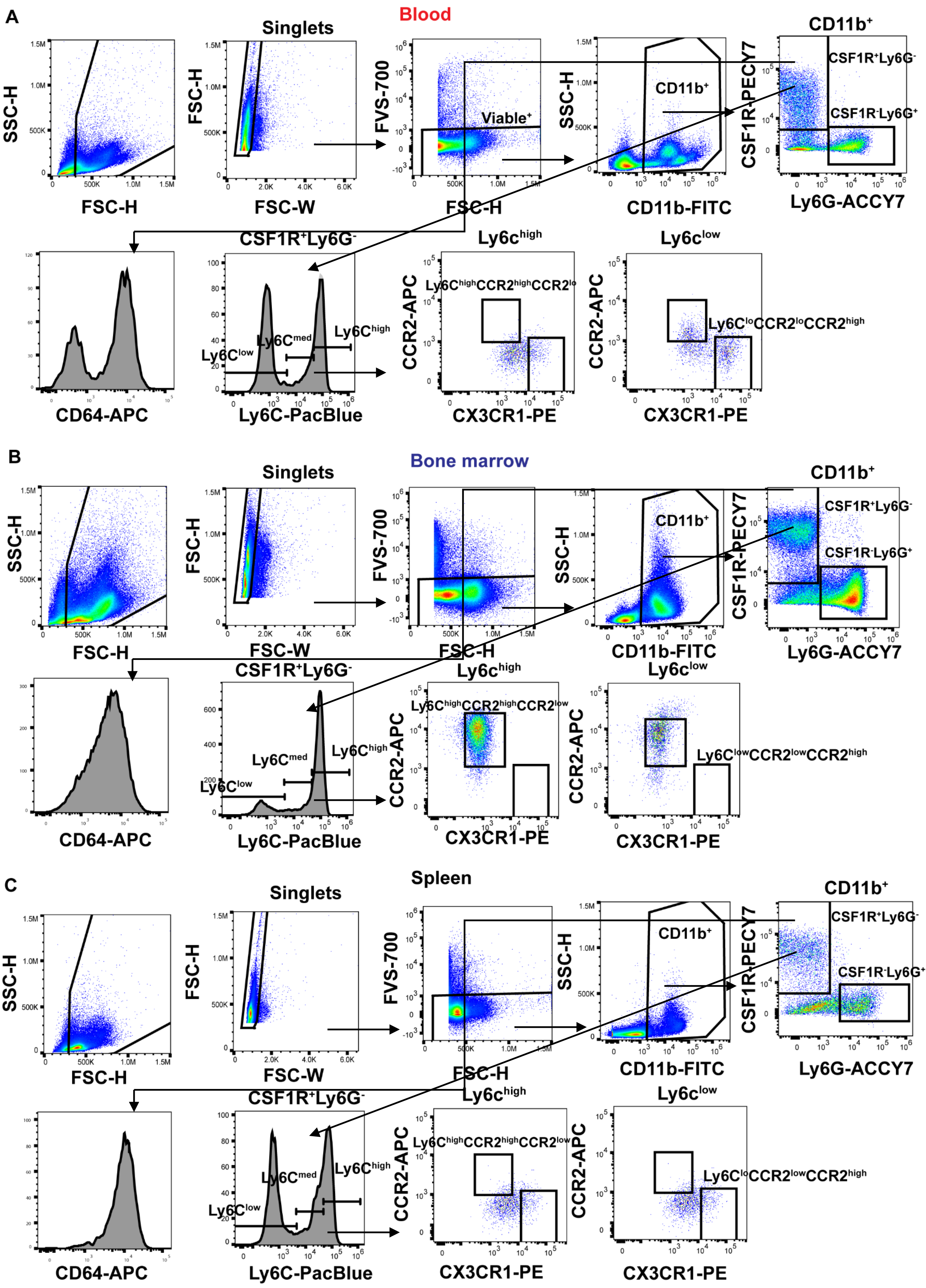

A
Blood
Singlets
SSC-H
FSC-H
FSC-W
FVS-700
Viable⁺
CD11b⁺
CD11b-FITC
CSF1R-PECY7
CSF1R⁺Ly6G⁻
CSF1R⁻Ly6G⁺
Ly6G-ACCY7
CSF1R⁺Ly6G⁻
CD64-APC
Ly6C^low
Ly6C^med
Ly6C^high
Ly6C-PacBlue
Ly6c^high
Ly6c^low
CCR2-APC
CX3CR1-PE
Ly6C^high CCR2^high CCR2^lo
Ly6C^lo CCR2^lo CCR2^high
B
Bone marrow
Singlets
SSC-H
FSC-H
FSC-W
FVS-700
CD11b⁺
CD11b-FITC
CSF1R-PECY7
CSF1R⁺Ly6G⁻
CSF1R⁻Ly6G⁺
Ly6G-ACCY7
CSF1R⁺Ly6G⁻
CD64-APC
Ly6C^low
Ly6C^med
Ly6C^high
Ly6C-PacBlue
Ly6c^high
Ly6c^low
CCR2-APC
CX3CR1-PE
Ly6C^high CCR2^high CCR2^low
Ly6C^low CCR2^low CCR2^high
C
Spleen
Singlets
SSC-H
FSC-H
FSC-W
FVS-700
CD11b⁺
CD11b-FITC
CSF1R-PECY7
CSF1R⁺Ly6G⁻
CSF1R⁻Ly6G⁺
Ly6G-ACCY7
CSF1R⁺Ly6G⁻
CD64-APC
Ly6C^low
Ly6C^med
Ly6C^high
Ly6C-PacBlue
Ly6c^high
Ly6c^low
CCR2-APC
CX3CR1-PE
Ly6C^high CCR2^high CCR2^low
Ly6C^lo CCR2^low CCR2^high

**Supplemental Figure 11. Gating strategy to identify different subsets of monocytes in the blood, BM and spleen. A-C)** Representative example is taken from the blood (A), BM (B) and spleen from a wildtype (C57BL/6) DSS-treated mouse. Singlet cells were gated according to forward and side scatter and then gated for viable (FVS-negative). Viable cells were then gated for $CD11b^{+}$ myeloid cells. $CD11b^{+}$ cells were gated to identify $CS1R1^{+}Ly6G^{-}$ monocytes or $CS1R1^{-}Ly6G^{+}$ granulocytes. $CD64^{+}$ monocytes were gated on $CD11b^{+}CS1R1^{+}Ly6G^{-}$ monocytes. Inflammatory monocytes were identified as $LY6C^{high}CCR2^{+}CX3CR1^{low}$ patrolling monocytes were identified as $LY6C^{lo}CCR2^{low}CX3CR1^{high}$.
